# Decoupling of interacting neuronal populations by time-shifted stimulation through spike-timing-dependent plasticity

**DOI:** 10.1101/2022.06.29.498110

**Authors:** Mojtaba Madadi Asl, Alireza Valizadeh, Peter A. Tass

## Abstract

The synaptic organization of the brain is constantly modified by activity-dependent synaptic plasticity. In several neurological disorders, abnormal neuronal activity and pathological synaptic connectivity may significantly impair normal brain function. Reorganization of neuronal circuits by therapeutic stimulation has the potential to restore normal brain dynamics. Increasing evidence suggests that the temporal stimulation pattern crucially determines the long-lasting therapeutic effects of stimulation. Here, we tested whether a specific pattern of brain stimulations can enable the suppression of pathologically strong inter-population synaptic connectivity through spike-timing-dependent plasticity (STDP). More specifically, we tested how introducing a time shift between stimuli delivered to two interacting populations of neurons can effectively decouple them. To that end, we first used a tractable model, i.e., two bidirectionally coupled leaky integrate-and-fire (LIF) neurons, to theoretically analyze the optimal range of stimulation frequency and time shift for decoupling. We then extended our results to two reciprocally connected neuronal populations (modules) where inter-population delayed connections were modified by STDP. As predicted by the theoretical results, appropriately time-shifted stimulation causes a decoupling of the two-module system through STDP, i.e., by unlearning pathologically strong synaptic interactions between the two populations. Based on the overall topology of the connections, the decoupling of the two modules, in turn, causes a desynchronization of the populations that outlasts the cessation of stimulation. Decoupling effects of the time-shifted stimulation can be realized by time-shifted burst stimulation as well as time-shifted continuous simulation. Our results provide insight into the further optimization of a variety of multichannel stimulation protocols aiming at a therapeutic reshaping of diseased brain networks.

## Introduction

Synaptic connections in cortical networks are highly adaptive due to activity-dependent synaptic plasticity [1]. Reshaping of connectivity patterns by plasticity mechanisms based on external stimuli is necessary for normal brain function such as appropriate motor actions [2]. Spike-timing-dependent plasticity (STDP) provides a mechanistic model for the modification of the strength of synaptic connections according to the temporal coincidence of pre- and post-synaptic activity [3–6]: Synapses are strengthened when the presynaptic spike precedes the postsynaptic spike, whereas they are weakened in the reverse scenario [4]. In this way, neuronal activity shapes the synaptic organization of brain networks [1, 7] which, in turn, adjusts the activity of neurons in a feedback loop [8–10]. Pathological conditions such as Parkinson’s disease (PD) [11–15], essential tremor [16–18] and epilepsy, [19–21] are connected with abnormal brain activity and connectivity.

In PD, for example, neurodegeneration triggers a cascade of compensatory or maladaptive changes within the cortico-subcortical circuits [22–24] that can lead to abnormal neuronal activity, e.g., excessive neuronal synchronization [11,12]. This can result in a reshaping of neuronal circuits due to the emergence of pathological connectivity patterns, e.g., strong synaptic connections [13, 14], through synaptic plasticity [24–26]. High-frequency (> 100 Hz) deep brain stimulation (HF-DBS) is an effective clinical therapy for medically refractory PD which can temporally relieve several motor symptoms of PD that are linked to abnormal neuronal synchronization [27, 28]. However, reappearance of symptoms soon after the discontinuation of stimulation [29] entails chronic stimulation which may further side effects [30, 31]. Also, DBS delivered to the subthalamic nucleus (STN) and globus pallidus internus (GPi) is considered as ineffective for treating impairment of gait and balance and is little beneficial for or even worsens speech impairment [32]. Based on computational studies, it was suggested to counteract abnormal neuronal synchrony by stimulation techniques designed to specifically cause desynchronization [33, 34].

In computational studies in oscillator and neuronal networks, different scenarios have been proposed for the desynchronization of neural populations by single-site stimulation (i.e., targeting one population) or multi-site stimulation (i.e., targeting two or more populations). For instance, desynchronization may be realized by demand-controlled delayed feedback stimulation [35–38] where the whole network is stimulated and registered at the same time which can be challenging in a clinical situation. This can be resolved by spatially splitting the whole population into two separate subpopulations, one being stimulated and the other being measured [39]. A more efficient approach has been suggested to separate the stimulation and registration processes in time rather than in space [40], i.e., by time-delayed feedback control of the pathological activity. However, these methods require a real-time measurement of the network activity. More importantly, smooth, non-pulsatile feedback stimulation techniques typically violate safety requirements, in particular, charge density limits [41–43]. Hence, based on computational studies, it was suggested to use linear and nonlinear delayed feedback signals to modulate amplitude and sign of continuous pulse train stimulation [44–46]. Alternatively, using multiple sites and tuning the temporal pattern of stimulation may have huge consequences on the stimulation outcome [47–49]. For instance, coordinated reset (CR) [50] stimulation is a theory-based multichannel patterned stimulation that targets subpopulation of neurons at different sites sequentially, i.e., in a timely coordinated manner [50, 51]. Computationally, CR stimulation can shift network dynamics from pathologically synchronized states to more physiologically favored states with desynchronized activity and, hence, induce long-lasting desynchronizing effects that outlast stimulation offset [51]. The desynchronizing effects [50], cumulative effects [52] and long-lasting effects [51] of the CR stimulation were validated in pre-clinical as well as clinical proof-of-concept studies [53–57].

Prolonged stimulation effects are desirable since they can induce sustained therapeutic outcome that outlast the cessation of stimulation. However, desynchronizing stimulation may not necessarily weaken pathologically strong synapses between the neurons during stimulation [58, 59]. Rather, long-lasting desynchronizing effects can be realized by decoupling neurons [59], i.e., by desynchronizing the overly synchronized activity of neurons and, furthermore, reduction of the pathologically strong synaptic connections between neurons. Overly synchronized neuronal activity can arise due to either strong local connections within the brain regions or due to the excessively potentiated long-range connections between the different brain regions [60–63]. Therefore, therapeutic stimulation techniques aiming at the reduction of abnormally synchronized activity could target both local connections and long-range projections.

In the normal brain, inter-areal communication and information exchange takes place through long-range synaptic connections [62–64]. However, pathological changes in inter-areal connectivity may impair normal brain function. For instance, PD can cause structural and functional reorganization of cortex characterized by altered inter-hemispheric neuronal activity and connectivity patterns [65,66]. Cortical stimulation is one of the therapeutic strategies that typically requires a less invasive surgical procedure in comparison to DBS surgery. Electrical stimulation of the motor cortex can lead to functional motor recovery in experimental models of PD [67,68]. Experimentally, it was shown that cortical stimulation can induce therapeutic changes in synaptic connectivity between two interacting networks [69–73], in this way restoring relevant features of physiological connectivity. As shown both computationally and experimentally [73,74], rewiring of synaptic connectivity between two interacting populations can be realized, e.g., by applying a specific set of stimuli to two populations to induce a time shift (delay) between their activity so that inter-population synapses could be regulated through STDP. This motivated us to study a stimulation paradigm applied to a cortical network model with more realistic assumptions where we explicitly considered transmission delays along with STDP. We hypothesized that appropriate temporal detuning of two stimulus trains delivered to two different populations, e.g., a time shift between stimuli, could effectively decouple the two populations through STDP and induced pronounced effects outlasting the cessation of stimulation. In fact, one of the goals of this computational study is to demonstrate how minor parameter changes in comparably simple stimulation protocols may massively change stimulation outcome.

To test our hypothesis, we used a generic cortical model to study theoretical conditions for decoupling two initially strongly coupled neuronal populations (modules) by repetitive stimulation of both populations with a time shift. To provide a theoretical basis for the simulation results, we first considered a reciprocally coupled two-neuron motif with plastic synapses and theoretically analyzed the optimal range of stimulation frequency and time shift to decouple neurons with a given set of STDP parameters and transmission delays. We then simulated a two-module model of cortical networks composed of weakly coupled excitatory and inhibitory neurons within each population. The two populations interacted via inter-population excitatory-to-excitatory delayed connections modified by STDP. In this study we focused on the modification of interpopulation synapses between the two modules to study the effect of the time-shifted two-site stimulation on the evolution of the plastic synapses between the two populations and the resultant effect on the dynamics of the network. To this end we assumed that the local connections within each population are weak and static. These assumptions meant that the synchronized activity of the network was due to the pathologically strong long-range connections and, furthermore, helped us to capture the pure effect of the modification of the long-range connections on the reduction of synchronized oscillatory activity of the network. Due to the initially strong inter-population synapses, the initial activity of the modules mimicked a pathological condition characterized by high firing rates of the neurons and large-amplitude collective oscillations. Stimuli were separately delivered to the two modules, each of them affecting an entire module, respectively. A dedicated time shift between the stimulus trains for the two modules modified inter-population synapses through STDP. We showed that the time-shifted stimulation enables the network to unlearn pathologically strong synaptic interactions between the two modules. Effective decoupling of the two modules ultimately caused a desynchronization of the network that persisted after stimulation cessation.

We showed that the time shift scheme may work in a rather generic manner. It can be realized by time-shifted trains of single stimulus pulses as well as patterned delivery of time-shifted bursts. The STDP potentiation and depression rates and time constants, and the delay in the transmission of signals along the inter-population connections determine the ultimate stimulation-induced distribution of the synaptic strengths after stimulation offset. Our generic results may contribute to the further development of temporally patterned stimulation in a variety of multichannel stimulation protocols optimized for unlearning pathological connectivity between neurons which can be adapted for cortical stimulation to induce long-lasting therapeutic effects by shifting the dynamics of the diseased brain towards healthy attractor states.

## Methods

### Neuron and Network Model

The pairwise analysis was performed on a two-neuron motif comprising two excitatory leaky integrate-and-fire (LIF) neurons [75] connected by reciprocal plastic synapses (Fig. 1A). As a model of cortical neuronal networks [60, 63, 76], two populations (modules), e.g., in two hemispheres (see Fig. 1B), were connected via inter-population plastic excitatory-to-excitatory synapses (Fig. 1C). Each module consists of *N* = 200 weakly coupled LIF neurons randomly connected in an sparse manner [77], of which *N*_ex_ = 0.8*N* were excitatory and *N*_in_ = 0.2*N* were inhibitory. In the normalized units, subthreshold dynamics of the dimensionless membrane potential (*v*_*i*_(*t*) = *V*_*i*_(*t*)*/V*_th_) of neuron *i* is described by the following differential equation:

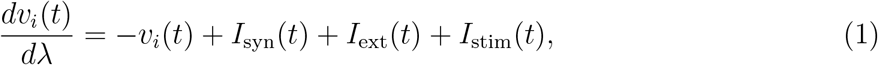

**Figure 1:**
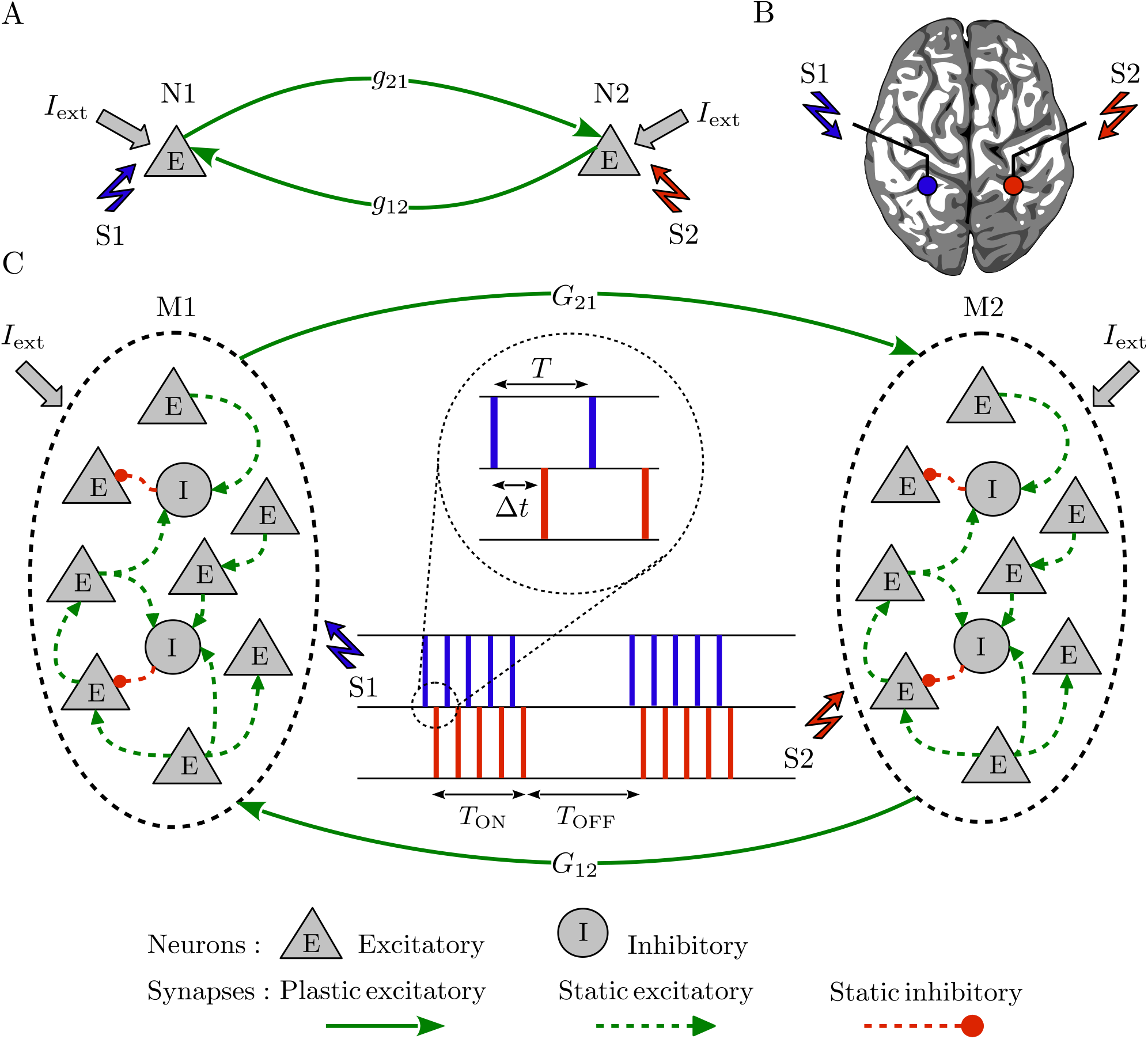
Representation of the two-neuron motif and neuronal network. (**A**) Two excitatory neurons coupled by reciprocal plastic synapses with strength *g*_21_/*g*_12_. (**B**) Schematic view of the brain and the stimulation electrodes (circles) where the two stimulation signals (S1/S2) were separately delivered to two representative populations in panel C, e.g., in two hemispheres. (**C**) Two reciprocally connected neuronal populations (modules) characterized by inter-population excitatory-to-excitatory plastic synapses with mean coupling *G*_21_/*G*_12_. Each stimulation signal consists of intermittent bursts where *T* represents inter-pulse interval within a burst and ∆*t* is the time shift between the two stimulation signals delivered to each module. *T*_ON_ is the stimulation ON-epoch for each stimulation burst and *T*_OFF_ is the stimulation OFF-epoch between two successive bursts within each stimulation signal.

where *λ* = *t/τ*_m_ is the dimensionless time in the units of the membrane time constant *τ*_m_ = 10 ms. When *v*_*i*_(*t*) reaches the firing threshold *v*_th_ = 1, the neuron fires and the membrane potential resets to the resting value *v*_r_ = 0. 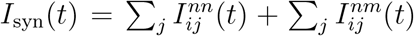 is the synaptic current, where 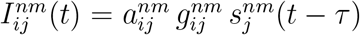 is the intra- (*n* = *m*) or inter-population (*n* ≠ *m*) synaptic current from the presynaptic neuron *j* in module *m* to the postsynaptic neuron *i* in module *n*. 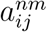 and 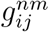 are the corresponding elements of the adjacency (**A**) and synaptic strength (**G**) matrices, respectively. *τ* = *τ*_d_ + *τ*_a_ is the inter-population delay in forward or backward direction, i.e., the sum of dendritic (*τ*_d_) and axonal (*τ*_a_) transmission delays in the synapse connecting the pre- and postsynaptic neurons. 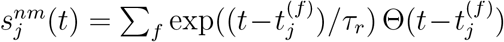 denotes the spiking activity of neuron *j* with a time constant *τ*_*r*_ = 5 ms [78], where *t*^(*f*)^ is the firing time of neurons and Θ(*t*) is the Heaviside step function. *I*_ext_(*t*) represents the spontaneous background activity due to the input from other brain areas modeled as a homogeneous Poisson process [60, 63].

### Stimulation Protocol

*I*_stim_(*t*) in Eq. (1) represents the stimulation current (S1/S2 in Fig. 1) composed of intermittent bursts that are simultaneously delivered to all (excitatory and inhibitory) neurons embedded within a population:

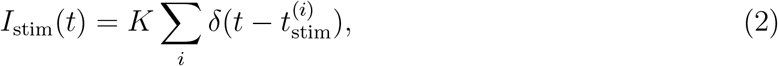

where *K* is a dimensionless parameter representing the stimulation intensity, 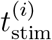 denotes the onset time of the individual stimulation pulses and *δ*(*t*) is the Dirac delta function. Given the initial point of the stimulation 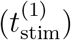, the time of the next pulse onset is determined by the following stimulation protocol:

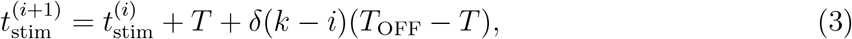

where *T* is the inter-pulse interval which alternatively represents the intra-burst frequency, i.e., *ν* = 1*/T*. The total duration of stimulation, i.e., *stimulation epoch*, was *T*_stim_ = 5 s. *k* = 5 is the number of pulses within a burst delivered for the duration of ON-epoch (*T*_ON_ = 120 ms). *T*_OFF_ = 360 ms represents the stimulation OFF-epoch between two successive stimulation bursts, i.e., the inverse of burst delivery rate, within each stimulation signal (see Fig. 1C).

S1/S2 stimulation signals could separately target two neurons (N1/N2 in Fig. 1A) or two representative populations (M1/M2 in Fig. 1C) in two hemispheres as shown in Fig. 1B. Assuming that the S1 stimulation signal is delivered to N1 (M1) at 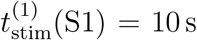, the S2 stimulation signal is delivered to N2 (M2) with a time shift 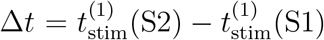 that represents the time shift between the onset times of stimulations delivered to the two neurons (populations), as measured between the first pulse in each train.

### Spike-Timing-Dependent Plasticity (STDP)

Pre- (*j*) and postsynaptic (*i*) neurons within each module were connected to each other via instantaneous static synapses with strength 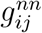, whereas the two modules were connected by inter-module excitatory-to-excitatory plastic synapses with transmission delays from the presynaptic neuron *j* in module *m* to the postsynaptic neuron *i* in module *n* with strength 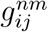 modified according to the following pair-based STDP rule [5]:

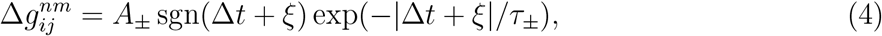

where *A*_*±*_ and *τ*_*±*_ are the learning rate and the effective time constant of synaptic potentiation (upper) and depression (lower sign), respectively, and sgn(∆*t*) is the sign function. ∆*t* = *t*_post_ − *t*_pre_ is the instantaneous time lag between pre- and postsynaptic spike pairs and *ξ* = *τ*_d_ − *τ*_a_ indicates the effective delay perceived at the synapse, i.e., the difference between dendritic and axonal transmission delays [79].

Evaluated over the time interval between two successive spikes (*T*), the potentiation and depression terms in Eq. (4) compete to determine the net synaptic change in a synapse as follows [79]:

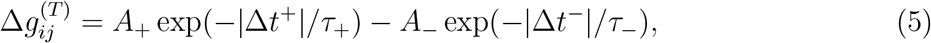

assuming that |∆*t*^+^| = ∆*t* + *ξ* is the time lag used by the STDP rule for potentiation of the synapse and |∆*t*^*−*^| = *T* − |∆*t* + *ξ*| is the depression time lag.

The synaptic strengths were updated by an additive rule at each step of the simulation, *g* → *g* + ∆*g*. The value of the synaptic strengths was restricted in the range [*g*_min_, *g*_max_] ∈ [0.05, 1]. The synaptic strengths were set to *g*_min_ (*g*_max_) via hard bound saturation constraint once they crossed the lower (upper) bound of their allowed range.

### Data Analysis

#### Inter-Population Mean Coupling

The mean coupling strength between the modules from the presynaptic module *m* to the post-synaptic module *n* was measured by calculating the average of the plastic inter-population synaptic strengths at a given time:

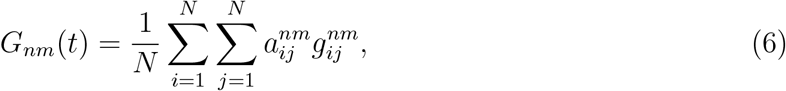

where *N* is the total number of neurons in each module. Furthermore, the time averaged inter-population mean coupling between the two modules in the network (*G*_ave_) was evaluated over 10 s of network activity after stimulation offset.

#### Population Activity

The population activity of module *i* in the network was calculated by counting the number of spikes in a time interval which gives the number of active neurons at that interval:

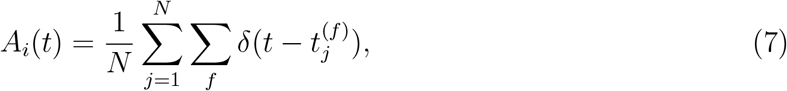

where *N* is the total number of neurons in module *i* and *t*^(*f*)^ is the firing time of individual neurons.

#### Pairwise Correlations

The Pearson correlation coefficient was used to calculate the spike count correlation between pairs of neurons (*i, j*) in each module that is given by:

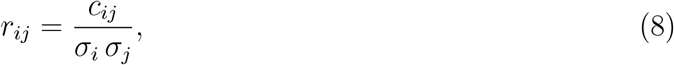

where *c*_*ij*_ is the covariance between spike counts of the two neurons calculated from their spike trains, and *σ*_*i*_ is the standard deviation of spike time distribution given by the corresponding spike train. Correlations were calculated based on the spike times resulted from 10 s of network activity before/after stimulation on/offset.

#### Spike Count Irregularity

The coefficient of variation of the inter-spike intervals (ISIs) was calculated as a measure of the irregularity of spiking activity of neuron *i*:

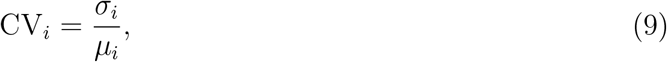

where *σ*_*i*_ is the standard deviation and *µ*_*i*_ is the mean of the ISIs calculated from the spike time distribution of neuron *i* given by its spike train evaluated over 10 s of network activity before/after stimulation on/offset.

#### Population Fano Factor

The population Fano factor (pFF) was used to measure the synchrony of population activity in module *i*, defined as [80]:

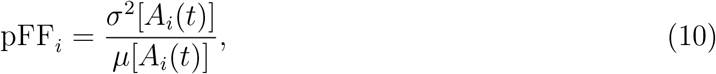

where *A*_*i*_(*t*) represents the population activity defined in Eq. (7), and *σ*^2^ and *µ* are the variance and mean of the population activity, respectively. The pFF evaluates the normalized amplitude of the variation of the population activity which increases when the neurons fire in synchrony [63, 81]. Smaller values of the pFF correspond to desynchronized states, whereas greater values of the pFF imply synchrony in the network. The pFF was evaluated over 10 s of network activity after stimulation offset.

## Results

### Two-Neuron Motif

We first studied if the time-shifted stimulation can decouple a two-neuron motif, i.e., reduce the initially strong synaptic connections and, furthermore, suppress the firing activity of neurons. For this, we explored the evolution of the synaptic strengths in a motif consisted of two stochastically firing excitatory neurons coupled by reciprocal plastic synapses (see Fig. 1A). This motif is representative of two neurons in two different populations connected by long-range connections and as will be shown below by demonstrating that results obtained in the motif can adequately predict those for two connected populations. For simplicity and for analytical tractability, we considered the classical STDP profile characterized by asymmetric modification of the synapses based on Eq. (4) [5]. We set the parameters of STDP qualitatively in accordance to the experimentally reported values for cortical regions [6], which is a generic form for the STDP profile characterized by larger potentiation rate (i.e., *A*_+_ *> A*_*−*_) and greater time constant of depression (i.e., *τ*_+_ *< τ*_*−*_).

We then stimulated the two neurons periodically with period *T* (or alternatively with frequency *ν* = 1*/T*) but at different times with a time shift ∆*t* in order to find which values of *T* and ∆*t* lead to a depression of both synapses. First, we ignored the delay in the transmission of the signals between the two neurons (i.e., when | *ξ* | = 0). Over a period each synapse experiences a succession of potentiation and depression and the net change of the synaptic strength depends on the superposition of the two changes [79, 82]. For more clarity we assumed that the neuron N1 in Fig. 1A is stimulated first and the neuron N2 is stimulated after the time shift ∆*t*. The 1 → 2 synapse experiences a potentiation 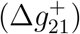 and a depression 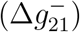, and it will be depressed if 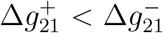. By the same token, the reverse synapse (i.e., 2 → 1) is depressed when 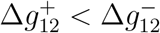. Taken together, the condition for depression of synapses in both directions will be given by 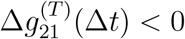 and 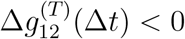 (see Eq. (5) in Methods).

The regions for depression (blue) and potentiation (red) of both synapses and the potentiation of one synapse and the depression of the other (orange) are shown Fig. 2 in the ∆*t*-*T* plane, as predicted theoretically by calculating the net synaptic change for reciprocal synapses between two neurons given by Eq. (5). The results indicate that for the chosen STDP parameters, for most values of ∆*t* and *T* the synapses will be potentiated in one direction and will be depressed in the other, leading to unidirectional connectivity. For the simultaneous depression of synapses in two directions, which is the target of our time-shifted stimulation approach, there is a desired range for ∆*t* and *T* where the neurons should fire almost in anti-phase, i.e., ∆*t ≈ T/*2 = 1*/*2*ν*. The numerical results shown in Fig. S1 (see Supplementary Material) for an exemplary set of parameters (marked by point b in Fig. 2A) verify the theoretical prediction, valid in the absence of axonal and dendritic delays.

**Figure 2:**
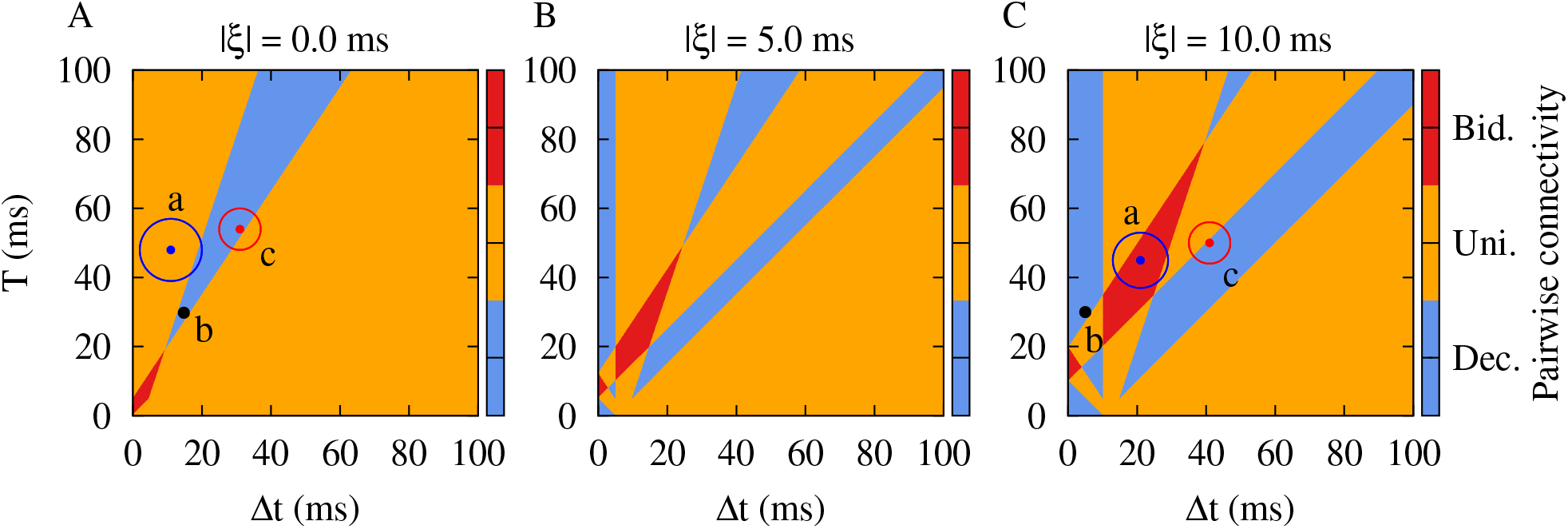
Theoretical prediction of the emergent two-neuron connectivity. Qualitative colors show the synaptic structure between a pair of pre- and postsynaptic neurons calculated based on the synaptic change in Eq. (5) over a period (*T*): Decoupled (blue), unidirectional (orange) and bidirectional (red) regimes. STDP parameters were *A*_+_ = 0.008, *A*_*−*_ = 0.005, *τ*_+_ = 10 ms and *τ*_*−*_ = 20 ms. (**A**) Points a: (11, 48) ± 9 ms, b: (15, 30) ms and c: (31, 54) ± 6 ms represent (∆*t, T*) pairs obtained from simulations performed in Fig. S1 before, during and after the stimulation, respectively, for | *ξ* | = 0.0 ms. (**B**) | *ξ* | = 5.0 ms; the effective delay at synapse reshapes the ∆*t*-*T* plane. (**C**) Points a: (21, 45) ± 8 ms, b: (5, 30) ms and c: (41, 50) ± 6 ms show the same (∆*t, T*) pairs as in A, but obtained from numerical simulations performed in Fig. 3 before, during and after the stimulation, respectively, for |*ξ*| = 10.0 ms.

Next, we explicitly considered the role of realistic transmission delays in the model. As shown previously [79, 82], the difference between the dendritic and axonal delays (*ξ* = *τ*_d_ − *τ*_a_), enters the formula for the modification of the synapses in Eq. (4), since the effect of the firing of pre- and postsynaptic neurons builds up at the synapse after axonal and dendritic (back-propagation) transmission delays, respectively. In this case, the condition for depression of the synapses in two directions is given by 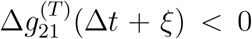 and 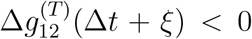. In Fig. 2B and C, regions for three different kinds of final pairwise synaptic connections are shown for two different values of | *ξ* |, representing median transmission delay between cortical populations [83]. We considered positive values of *ξ* since for long-range connections axonal delays are greater than dendritic delays. Intriguingly, at small values of the firing period (i.e., at high stimulation frequencies), and also at greater periods (i.e., at low stimulation frequencies), a close-to in-phase firing may lead to depression of both synapses. This is illustrated by a numerical experiment in Fig. 3A and B with exemplary parameters marked by point b: The simulation related to (∆*t, T*) = (5, 30) ms in Fig. 2C demonstrates the validity of the analytical predictions. Note, with this set of parameters coincidence of the spontaneous firing of the neurons led to a potentiation of both synapses (point a in Fig. 2C). In contrast, stimulation with appropriate parameters (point b in Fig. 2C) led to a depression of both synapses after stimulation offset (point c in Fig. 2C). Obviously, administration of additional trains of stimuli leads to a further depression of the synapses. Fig. 3C also shows that in the two-neuron motif either with (bottom) or without (top) transmission delay, the time-shifted stimulation effectively reduced the mean firing rates in both cases. This lead to a desynchronization of the two-neuron motif by reducing the coincidence of the neuronal discharges where the neurons were unable to resynchronize their activity due to the weakened coupling.

**Figure 3:**
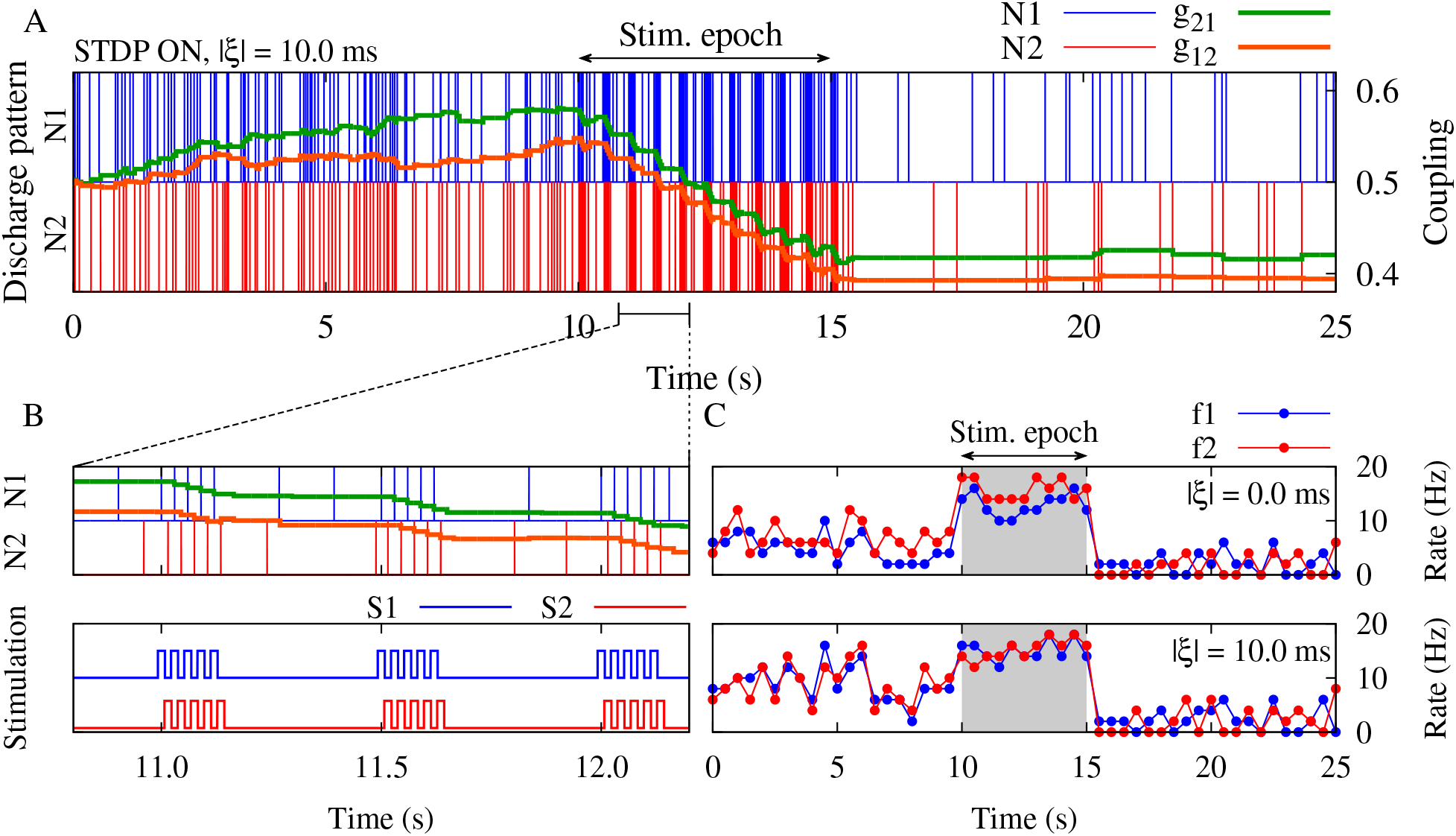
Suppression of the synaptic strengths between two neurons by time-shifted stimulation. (**A**) Time course of neuron discharges (N1/N2) and the synaptic strengths (*g*_21_/*g*_12_) are shown for two neurons when STDP parameters were *A*_+_ = 0.008, *A*_*−*_ = 0.005, *τ*_+_ = 10 ms and *τ*_*−*_ = 20 ms, and | *ξ* | = 10.0 ms. (**B**) (Top) Results in panel A are shown with a higher resolution. (Bottom) Stimulation signals were delivered to N1 (S1) and N2 (S2) for the duration of *T*_stim_ = 5 s (stimulation ON period in A) with time shift ∆*t* = 5 ms and frequency *ν* = 1*/*30 Hz (inspired by parameters shown in point b in Fig. 2C). (**C**) Time course of the mean firing rate of neurons (*f* 1*/f* 2) for |*ξ*| = 0.0 ms (top) and |*ξ*| = 10.0 ms (bottom).

### Bidirectionally Connected Populations

We then replaced each neuron in the two-neuron motif by a population of excitatory and inhibitory neurons and studied whether the theoretical predictions obtained in the two-neuron motif were valid for two bidirectionally connected populations (i.e., two modules in Fig. 1C). The approach employing two-neuron motifs and then translating the results to two interacting populations was successfully used previously in several computational studies [84–86]. By the same token, several studies used neuronal network models comprising two populations interacting via long-range excitatory-to-excitatory delayed projections modified by STDP [63, 64]. Weakly coupled neurons within each population were connected in a sparse manner and the individual neurons fired stochastically. Initially strong synaptic connections between the two populations led to synchronous firing of the two modules with high firing rate which is a marker of pathological conditions, e.g., in PD [12, 87]. Stimuli were delivered to the two populations separately, thereby homogeneously affecting all excitatory and inhibitory neurons in each population, respectively. By stimulating both populations with time-shifted stimulus pulse trains, we aimed at inter-population decoupling, i.e., reduction of the strong, plastic synaptic connections between the two populations, in this way inducing an effect outlasting the cessation of stimulation. This, in turn, caused a desynchronization of the populations. The parameters of the stimulation, including frequency and time shift, were chosen based on our results obtained in a pair of neurons as shown in Fig. 2.

We first examined the system in the absence of any transmission delay. In that case, the stimulation can suppress the inter-population connections and desynchronize the neuronal activity in the populations in a long-lasting manner, exceeding cessation of stimulation (Fig. S2, Supplementary Material). Note, taking into account realistic transmission delays changes the stimulation parameters required for decoupling and desynchronizing the two cortical modules (see Fig. 2C). Experimental estimates of delays in long-range inter-population connections vary from a few to tens of milliseconds [88] and, hence, cannot be ignored in biologically realistic simulations. As shown previously [89, 90], in the absence of transmission delays synapses tend to evolve in an asymmetric manner, giving rise to unidirectional connections, where depression of a synapse usually comes at the expense of potentiation of the reverse synapse. Therefore, bidirectional depression of synaptic strengths is unlikely to occur in the absence of transmission delays, especially for balanced STDP profiles [89, 90]. In contrast, in the presence of transmission delays, simultaneous bidirectional depression or potentiation of reciprocal synapses is more likely to occur (see Refs. [79, 82, 91] and cf. Fig. 2A with B and C).

In this study, we fixed the dendritic (back-propagation) delay at a biologically realistic value *τ*_d_ = 0.5 ms, whereas the remaining delays were assigned to the axonal delay. The dynamical and structural characteristics of the network before and after stimulation are shown in Fig. 4. The initial values of the connection strengths were chosen such that the neurons in the two populations fired synchronously, while the two populations oscillated in an anti-phase manner (Fig. 4A1). Other phase relationships could be obtained depending on the choice of the inter-populations delays (see Fig. S2, Supplementary Material). This models of a pathological state with strong phase-locked oscillations of the two populations (Fig. 4D, left), regular spiking (Fig. 4C, grey) at a high rate (Fig. 4H) and high pairwise correlation (Fig. 4B, grey) between the spiking activities of the neurons in each module. Based on the values of the periods and the time-shift between the stimulus trains in the two-neuron motif (Fig. 2), the outcome of stimulation can be predicted. The simulation results for the two populations (Fig. 4F and G) are in accordance with the prediction of the theoretical results (Fig. 2C). To induce an unlearning of the pathological state, we choose stimulation parameters from the two-neuron motif given by point b in Fig. 2C in order to decouple the two populations (frequency *ν* = 1*/T* = 1*/*30 Hz, time shift ∆*t* = 5 ms and delay | *ξ* | = 10.0 ms). After 5 s of stimulation, the mean synaptic strengths between both cortical modules were significantly suppressed in both directions (Fig. 4F). Accordingly, right after stimulation the initially pronounced oscillations of both cortical modules are suppressed (Fig. 4A2 and E, colored), and the neurons fire at a much lower rate and in an irregular manner (Fig. 4H, B and C, colored).

**Figure 4:**
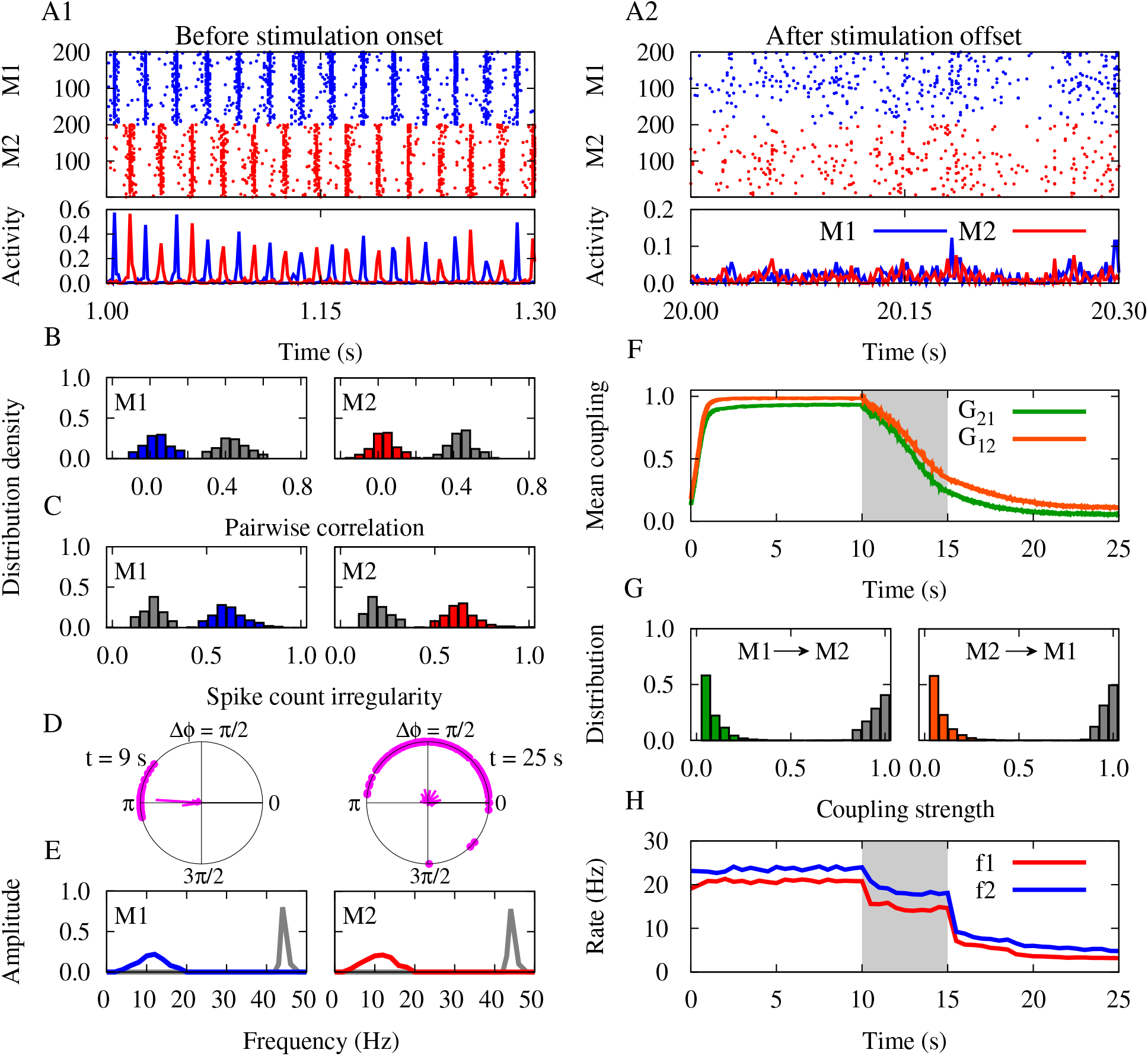
Decoupling by time-shifted stimulation in the network. (**A1**,**A2**) Raster plots and population activities (percentage of neurons firing per time window) are shown for the modules (M1/M2) before/after stimulation on/offset. (**B**,**C**) Distribution of the pairwise correlation and spike count irregularity of each module before (grey) and after (colored) stimulation epoch. (**D**) Snapshot of phase lags (∆*φ*) between synchronous discharges from the two modules before (*t* = 9 s, left) and after (*t* = 25 s, right) stimulation epoch. The radial bars show the distribution of the phase lags. (**E**) Fourier transform frequency of the population activity of each module before (grey) and after (colored) stimulation epoch. (**F**) Time course of the inter-population mean coupling (*G*_21_/*G*_12_). (**G**) Distribution of the inter-population synaptic strengths before (grey) and after (colored) stimulation epoch. (**H**) Time course of the mean firing rates (*f* 1*/f* 2) of neurons in each module. The modules were stimulated with time shift ∆*t* = 5 ms and frequency *ν* = 1*/*30 Hz (point b in Fig. 2C) for the duration of *T*_stim_ = 5 s (highlighted area in F and H). The effective delay was |*ξ*| = 10.0 ms. STDP parameters were *A*_+_ = 0.008, *A*_*−*_ = 0.005, *τ*_+_ = 10 ms and *τ*_*−*_ = 20 ms.

This realizes the decoupling and desynchronizing effects of the stimulation. In fact, stimulation of both populations with a time shift results in a significant reduction of inter-population coupling strength through STDP. This causes a desynchronization of the populations due to a decay of the external drive from the reciprocal population [92]. In this way, the decoupled populations return to their desynchronized firing activity in the absence of a significant external drive where the individual neurons fired stochastically. These irregular activity states reflect basic properties of normal cortical states [93, 94].

### Long-Lasting Decoupling by STDP

Intriguingly, after stimulation offset the connections continue to weaken. This is a very important point for the design of stimulation protocols that enable long-lasting effects. To avoid a relapse of the pathological dynamics the system has to be shifted into the basin of attraction of the desynchronized state, rather than still remaining in regimes supporting repotentiation of the synapses (i.e., red or orange zone in Fig. 2). In fact, the targets of our stimulus protocol are the basins of attraction of favorable, desynchronized states (cf. Ref. [95]) with STDP profiles satisfying the condition *A*_+_*τ*_+_ *< A*_*−*_*τ*_*−*_ [96] (blue region in Fig. 5A1/A2), where the irregular neuronal firing and the low inter-module correlation may enable a continued depression of the synapses between both cortical modules. We hypothesized that the decrease of the synaptic weights below a certain threshold can prevent from re-potentiation of the synapses and the corresponding relapse of pathological dynamics after the cessation of stimulation.

**Figure 5:**
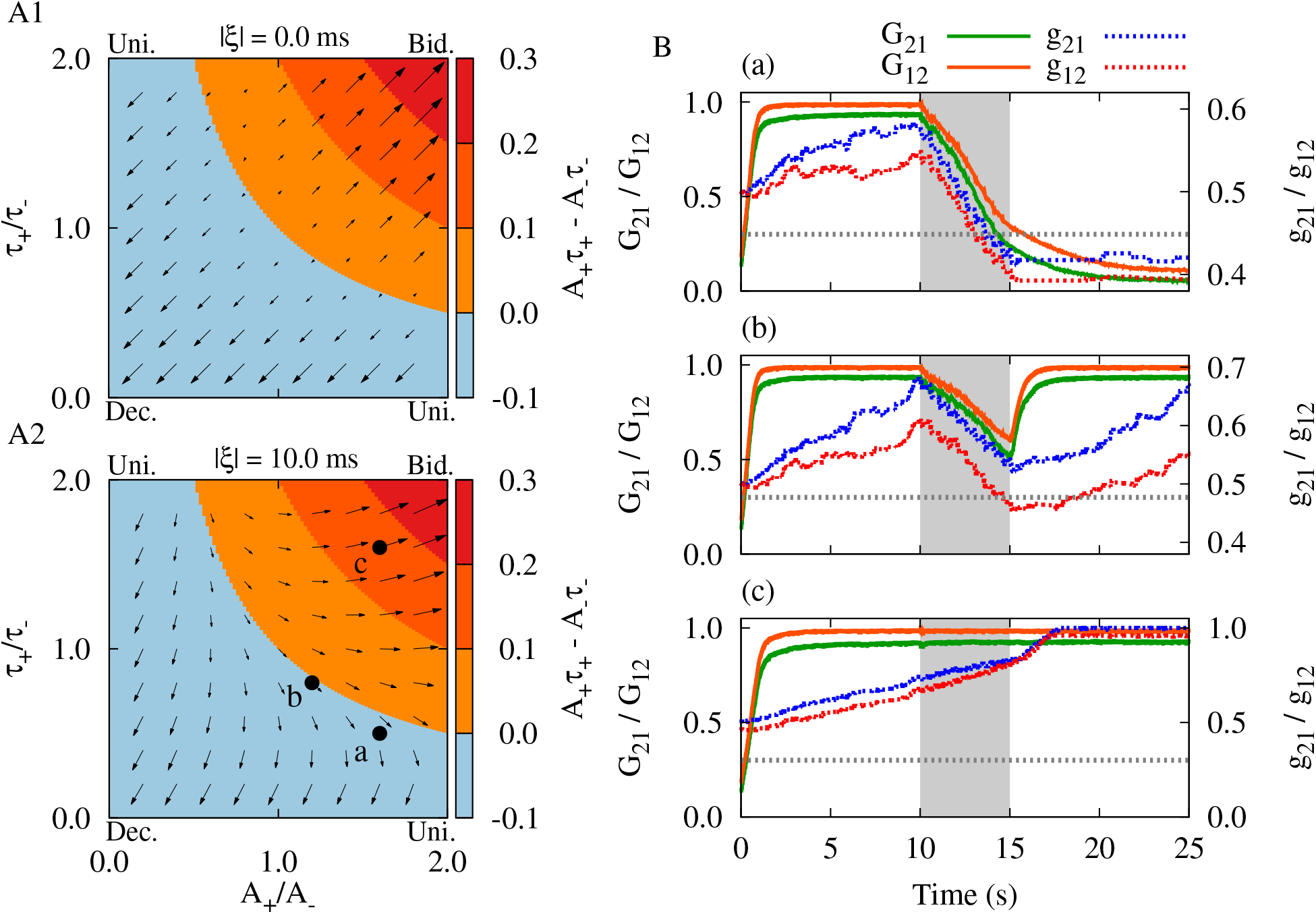
Stimulation-induced decoupling depends on the STDP parameters. (**A1**,**A2**) Colors show the imbalance between STDP potentiation and depression rates, and time constants for |*ξ*| = 0.0 ms (A1) and |*ξ*| = 10.0 ms (A2). *A*_*−*_ = 0.005 and *τ*_*−*_ = 20 ms were fixed and *A*_+_ and *τ*_+_ were varied. Arrows show the direction of synaptic change calculated theoretically based on Eq. (5) where each corner of A1 and A2 labeled as the emergent synaptic structure between two neurons marked as bidirectional (right top), unidirectional (left top and right bottom) and decoupled (left bottom) states. Point a: (*A*_+_*/A*_*−*_, *τ*_+_*/τ*_*−*_) = (1.6, 0.5) shows parameters used for stimulation in Figs. 3 and 4. Parameters shown in points b: (1.2, 0.8) and c: (1.6, 1.6) were used to provide a comparison. (**B**) Time course of the synaptic strengths in the two-neuron motif (*g*_21_/*g*_12_) and inter-population mean coupling between two modules in the network (*G*_21_/*G*_12_) are shown for STDP parameters represented by points a-c in A2, respectively. The highlighted area indicates the stimulation epoch. The horizontal dotted lines (grey) roughly show the threshold (*G*_th_ 0.3) where the synaptic strengths must cross to obtain a stable loosely connected network structure. Stimulation parameters were *ν* = 1*/*30 Hz and ∆*t* = 5 ms.

To test this hypothesis and to demonstrate the predictive ability of the two-neuron results we repeated the simulations for both the two-neuron motif and the bidirectionally connected neuronal populations with different sets of STDP parameters (three sets are shown in Fig. 5A2). Changing the ratio of potentiation and depression rates and time constants leads to different trade-offs between potentiation and depression regimes. For illustration, we used the same stimulation protocol with fixed parameters (*ν* = 1*/*30 Hz and ∆*t* = 5 ms) and a given delay | *ξ |* = 10 ms, and observed qualitatively different outcomes by changing the STDP parameters: Simultaneous depression of the synapses (Fig. 5B, a and b) or simultaneous potentiation of the synapses (Fig. 5B, c). While *A*_+_*τ*_+_ *< A*_*−*_*τ*_*−*_ was ensured in Fig. 5B (a and b), the mean rates of the depression of the synapses were different because of the different STDP parameters.

Whether or not the strong synaptic connections between the two neurons or two populations are weakened during stimulation and continue to weaken after the cessation of stimulation depends on the STDP parameters. The depression of the synaptic strengths below a threshold leads to a continued weakening of synaptic connections outlasting stimulation offset. (Fig. 5B, a). By changing the STDP parameters, it might take longer to enter the targeted basin of attraction. For instance, if at stimulation offset not all synaptic weights were below threshold, the connections got re-potentiated thereafter (Fig. 5B, b). In contrast, in Fig. 5B (c) the stimulation was unsuccessful since for the selected set of STDP parameters the stimulation induced a potentiation of the synapses. These behaviors were fairly predicted by the direction of the synaptic change (indicated by arrows) in Fig. 5A1 and A2 calculated from Eq. (5) for the two-neuron motif where each corner in panels A1 and A2 marks the emergent pairwise synaptic structure. A successful, long-lasting decoupling stimulation requires the stimulation frequency and time shift to be chosen in a way that STDP depression parameters dominate over potentiation. In addition, a sufficient amount of stimulation has to be delivered to ensure that the synaptic weights fall below threshold in order to cause a desynchronization of the populations.

### Burst Stimulation vs. Continuous Stimulation

So far, we used a time-shifted burst stimulation pattern (see Fig. 6A1, top) in order to suppress the synaptic connections between the two neurons (Fig. 3A) or the two modules (Fig. 4F). In this case, the depression of the synaptic strengths at the end of each stimulation epoch occurs in a step-like manner due to the ON/OFF epochs of the burst stimulation (*T*_ON_/*T*_OFF_ in Fig. 6A1, top). When the stimulation parameters, i.e., time shift, frequency and integral amount of stimulation, are adequately tuned based on the given STDP parameters, stimulation of sufficient duration can effectively decouple strongly connected neurons in the two-neuron motif as well as the two-module network.

**Figure 6:**
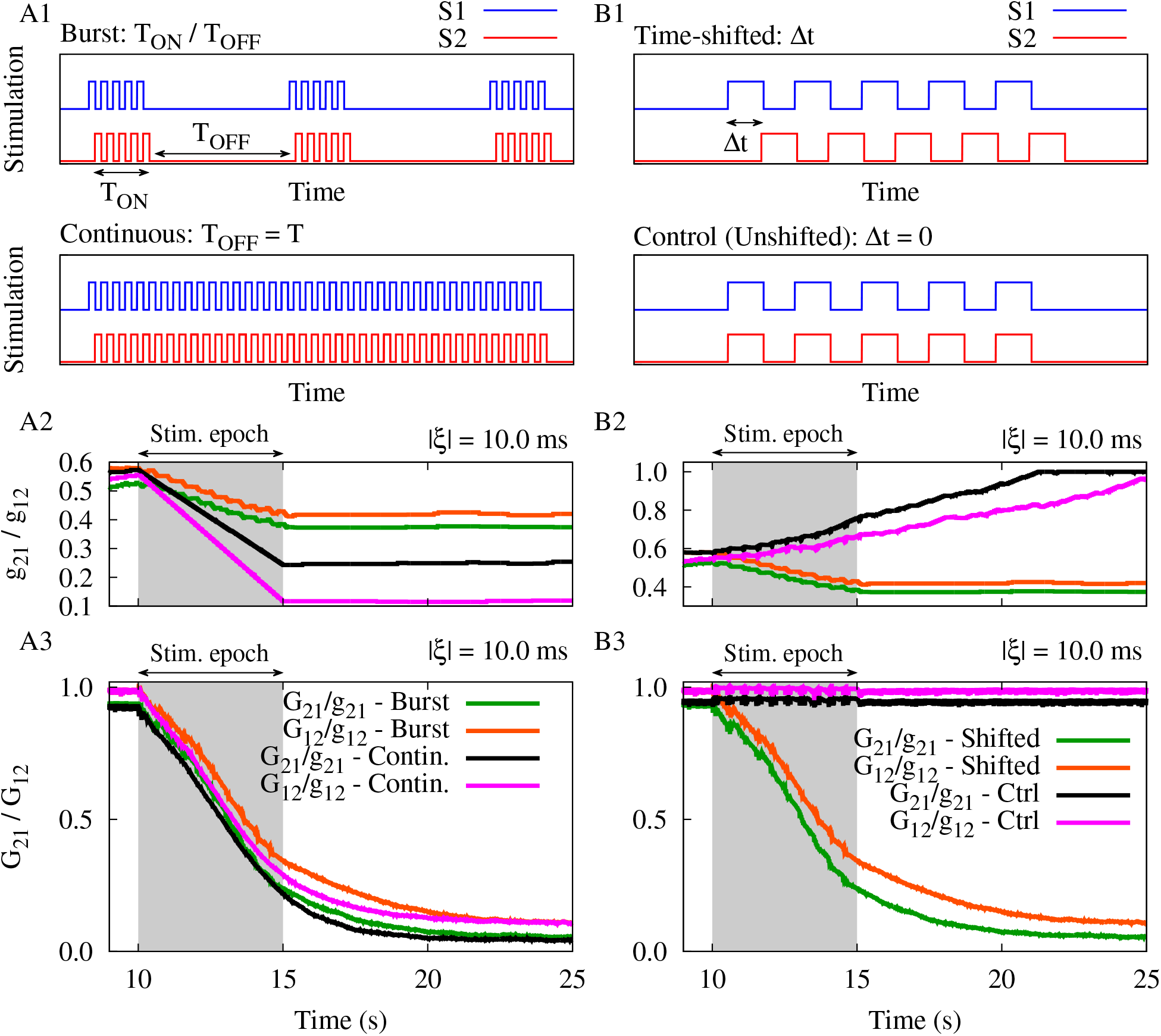
The effect of stimulation pattern on the stimulation outcome. (**A1**) Burst stimulation pattern (top) vs. continuous stimulation pattern (bottom) with the same frequency and time shift between the two signals (i.e., *ν* = 1*/*30 Hz and ∆*t* = 5 ms). (**A2**,**A3**) Time course of the synaptic strengths in the two-neuron motif (*g*_21_/*g*_12_) and the inter-population mean coupling between two modules in the network (*G*_21_/*G*_12_) are shown for burst and continuous patterns delivered to each module for | *ξ* | = 10.0 ms. (**B1**) Time-shifted stimulation (top) vs. control coincident stimulation (bottom) with the same frequency (*ν* = 1*/*30 Hz). (**B2**,**B3**) Same as A2 and A3, but for the burst stimuli delivered with and without time shift. The highlighted area indicates the stimulation epoch. STDP parameters were *A*_+_ = 0.008, *A*_*−*_ = 0.005, *τ*_+_ = 10 ms and *τ*_*−*_ = 20 ms.

To demonstrate whether mutually shifting the stimulation pulse patterns in time by ∆*t* may work in a rather generic way, we additionally considered a continuous stimulation pattern (Fig. 6A1, bottom), implemented by eliminating the OFF-epoch (*T*_OFF_) from the stimulation protocol by setting *T*_OFF_ = *T* in Eq. (3). Note that the continuous stimulation was applied to the system with the same intra-burst frequency (i.e., *ν* = 1*/*30 Hz) as of the burst stimulation. In both scenarios, the stimulation signals delivered to the two neurons or two modules were time-shifted by ∆*t* = 5 ms. As one can hypothesize, the application of a continuous stimulation pattern in this setting expedited the rate of decoupling due to the elimination of *T*_OFF_ (a time window enabling the synaptic strengths to re-increase) as it is shown in Fig. 6A2 and A3. However, the integral current delivery is much higher in the case of continuous stimulation. The results obtained with continuous stimulation indicate that the synaptic weight reduction induced by time-shifted stimulus delivery may be a mechanism that applies to a larger class of stimulation patterns, not only to periodic delivery of bursts.

### Time-Shifted Stimulation vs. Unshifted Stimulation

We illustrated that by introducing a simple time shift between the stimulation signals applied to two neurons or to two bidirectionally connected neuronal populations effectively reduced strong synaptic connections between the populations and caused a desynchronization of the populations. In particular, to demonstrate the significance of the time shift in the burst stimulation pattern (shown in Fig. 6B1, top) we considered a control condition, where bursts were delivered coincidentally, without time shift (shown in Fig. 6B1, bottom) and studied its impact on the two-neuron motif and the two-module network.

We repeated the simulations for the two-neuron motif and the two-module network, thereby taking into account realistic transmission delays in the model. The control condition was implemented by setting ∆*t* = 0. In this case, spontaneous background activity may cause jitters between in-phase stimulation pulses resulting in the emergence of small time lags between neuronal spikes which can lead to the potentiation of the synapses in both directions since *A*_+_ *> A*_*−*_ in Eq. (5). Therefore, in this setting the type of synaptic modification is determined by the dominance of the STDP-induced potentiation over STDP-induced depression rather than the STDP time constants (*τ*_+_ *< τ*_*−*_) that are important at greater time lags. As shown in Fig. 6B2 and B3, the control stimulation induced a reciprocal potentiation of the synapses between the two neurons in the two-neuron motif and, by the same token, potentiated the inter-population synaptic connections between the two bidirectionally connected neuronal populations (cf. green/orange curve with black/magenta one in Fig. 6B2 and B3).

### Rescaling Stimulation Frequency and Time Shift

In order to study the influence of the stimulation time shift (∆*t*) and frequency (*ν*) on the stimulation outcome we ran numerical simulations for a reasonable range of ∆*t* and *ν* and measured the averaged inter-population mean coupling between the two neuronal populations given by Eq. (6) and the synchrony level by the pFF of the activity of each module given by Eq. (10), as it is shown in Fig. 7. The pFF evaluates the normalized amplitude of the variation of the population activity which increases when the neurons fire in synchrony [63, 81].

**Figure 7:**
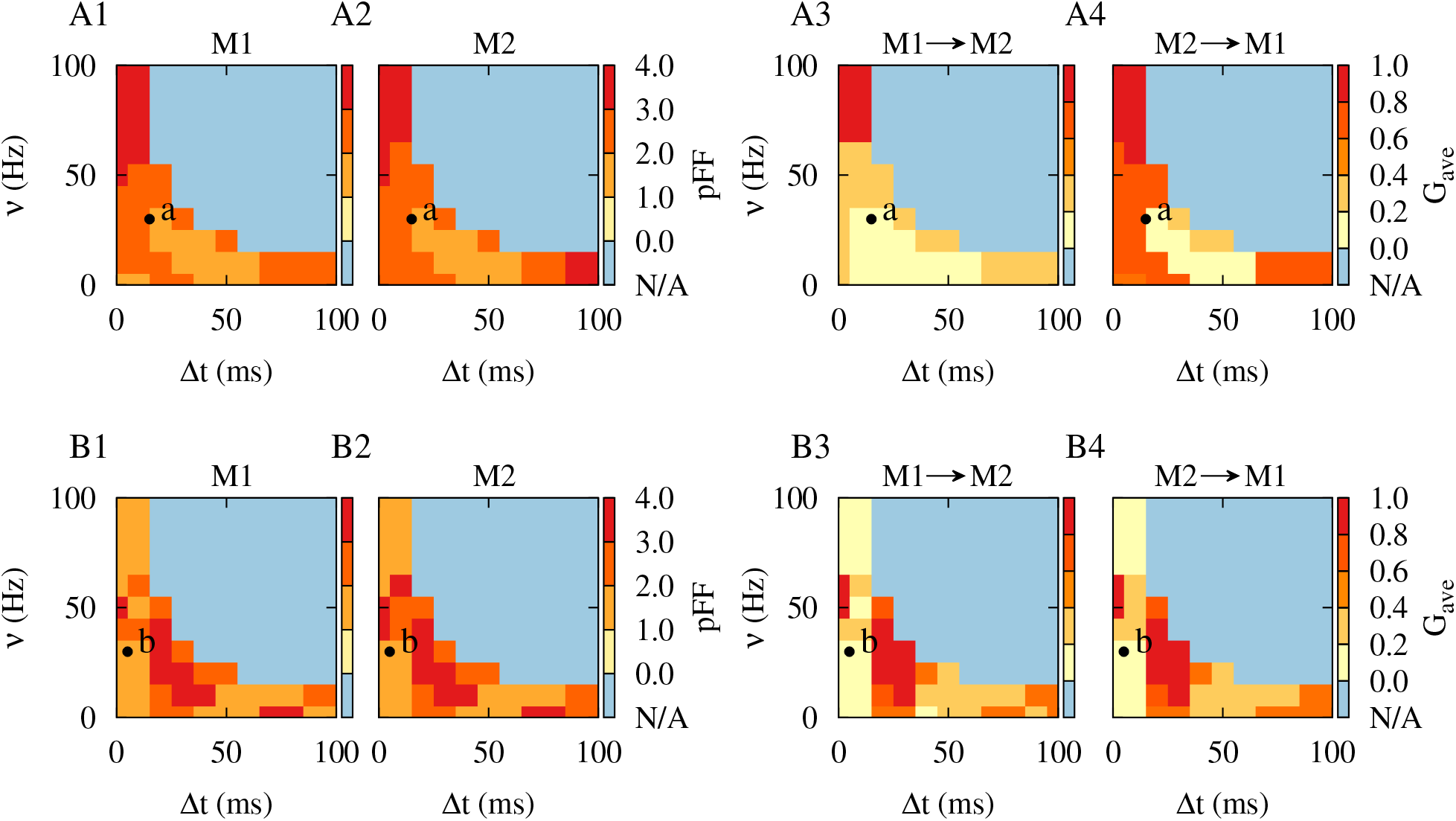
Rescaling the parameters of time-shifted decoupling stimulation. (**A1-A4**) Synchrony level measured by the pFF of the activity of each module (A1,A2) and the averaged inter-population mean coupling (*G*_ave_) between the two modules in the network (A3,A4) obtained by numerical simulations for different values of stimulation frequency and time shift with |*ξ*| = 0.0 ms, evaluated over 10 s of activity after stimulation offset. (**B1-B4**) Same as A1-A4, but with |*ξ*| = 10.0 ms. STDP parameters were *A*_+_ = 0.008, *A*_*−*_ = 0.005, *τ*_+_ = 10 ms and *τ*_*−*_ = 20 ms. Points a and b in the figure indicate the stimulation frequency and time shift used in Fig. 2A and C, respectively. Blue (N/A) region shows the range of parameters where the 0 *<* ∆*t < T* = 1*/ν* constraint is not satisfied.

When transmission delays were not considered in the model (Fig. 7A1-A4), stimulation frequencies over 50 Hz with a small time shift (i.e., ∆*t <* 10 ms) and stimulation frequencies below 20 Hz with a large time shift (i.e., ∆*t >* 70 ms) failed to desynchronize neuronal activity in each module (Fig. 7A1 and A2, dark red) or suppress strong inter-population connections between the modules (Fig. 7A3 and A4, dark red). However, stimulation frequencies below 50 Hz with greater time shifts (i.e., 10 ms *<* ∆*t <* 70 ms) induced desynchronizing effects (Fig. 7A1 and A2, light red) and suppressed inter-population connectivity in the network (Fig. 7A3 and A4, light red). Point a characterized by (∆*t, ν*) = (15 ms, 1*/*30 Hz) in Fig. 7A1-A4 shows the set of time shift and stimulation frequency used for decoupling stimulation in the absence of delays (see Figs. S1 and S2 in Supplementary Material). The blue (N/A) region in Fig. 7 shows the range of parameters where the constraint 0 *<* ∆*t < T* = 1*/ν* is not satisfied.

Inclusion of realistic transmission delays in the model crucially reshapes the optimal range of stimulation parameters required to induce decoupling effects (shown in Fig. 7B1-B4). In particular, in this case, stimulation frequencies over 60 Hz or below 30 Hz both with a small time shift (i.e., ∆*t <* 10 ms = |*ξ*|) induced desynchronizing effects (Fig. 7B1 and B2, light red) and suppressed inter-population connectivity (Fig. 7B3 and B4, light red). This was also seen for stimulation frequencies below 20 Hz with medium range time shifts (i.e., 40 ms *<* ∆*t <* 80 ms). However, intermediate stimulation frequencies (i.e., 10 Hz *< ν <* 50 Hz) with time shifts in the range 20 ms *<* ∆*t <* 30 ms can be detrimental since they induce synchronized activity states with strong inter-population synaptic connections (Fig. 7B1-B4, dark red). Point b, given by (∆*t, ν*) = (5 ms, 1*/*30 Hz), in Fig. 7B1-B4 corresponds to the time shift and stimulation frequency selected for decoupling stimulation in the presence of delays (see Figs. 3 and 4).

The theoretically predicted multistability of decoupled (blue), unidirectional (orange) and bidirectional (red) connectivity regimes for the two-neuron motif in Fig. 2 supports the stimulation-induced neuronal activity and synaptic connectivity emerging in the two-module network in Fig. 7 fairly well. The inventory of these attractor states allows to choose the parameters of the time-shifted stimulation appropriately so that the stimulation pattern can lead to an unlearning of pathologically strong synaptic connectivity between the two modules and cause a desynchronization of the modules (light red in Fig. 7), in this way inducing a sustained decoupling effect.

## Discussion

Our results illustrate that subtle changes of a stimulation protocol may have significant impact on the stimulation outcome. We presented a time-shifted two-channel stimulation approach in a generic cortex model that aims at decoupling two interacting neuronal populations by employing STDP, i.e., by unlearning pathologically strong synaptic interactions between the two populations. This simple intervention caused pronounced changes of the network dynamics and connectivity, outlasting the cessation of stimulation. In our model intra-population synaptic connections were weak and static and they had no role in the generation of synchronized oscillations and instead, synchronization of the populations was due to initially strong inter-population connections. Accordingly, decoupling of the two populations by the stimulation-induced reduction of inter-population excitatory projections below a threshold caused desynchronized and sparse activity of the neurons which is hallmark of the normal activity of cortical neurons [93,94]. Remarkably, this state was stable and the pathological connectivity and dynamics did not relapse after cessation of the external stimulus.

To reduce stimulation-induced side effects (see e.g. Ref. [31]) and to restore physiological function, it is desirable to reduce the integral amount and duration of stimulation. Accordingly, we do not only focus on acute effects (during stimulus delivery), but also on beneficial long-lasting effects, persisting after the cessation of stimulation [51, 52, 95]. Computationally, desynchronizing stimulation can shift adaptive networks from pathological attractor states towards physiological states [51,59,97]. In this way, therapeutic stimulation effects can be achieved that persist after discontinuation of stimulation [51, 59], as demonstrated in pre-clinical as well as clinical proof-of-concept studies [53–57]. However, the efficacy of desynchronizing stimulation crucially depends on the adaptation of stimulation parameters to the synchronization properties of the targeted network activity [59]. Furthermore, acute desynchronization does not necessarily lead to long-lasting changes of network activity [58, 59]. Apparently, enduring desynchronization can only be achieved if the pathological network connectivity is also modified by the external stimuli [58, 59], i.e., when the reduction of strongly synchronized activity is accompanied by a reduction of pathologically strong synaptic connectivity, to avoid relapse and reappearance of symptoms. Control of synchronization in neuronal populations and rewiring of synaptic connections by stimulation was previously addressed in a number of computational studies. For example, different patterns of sequential activations of subpopulations of neurons with a small intra-population delay (time shift) between multiple stimulation sites (see Fig. 8A) can induce an unlearning of pathological connectivity due to the desynchronization of neuronal activity in adaptive network models of phase oscillators [51] as well as a various neuron models [49, 95, 97–100]. Another study focused on harnessing the underlying STDP in the network to reshape synaptic connectivity by applying dedicated stimuli in a two-population network model of excitatory and inhibitory LIF neurons [86]. Anti-phase delivery of charge-balanced stimulation pulses to the excitatory subpopulation and the inhibitory subpopulation with a small time shift led to a reduction of the average synaptic weight between the two subpopulations in the network along with a strong desynchronizeation [86]. More recently, it was shown that two-site stimulation of cortical populations with long inter-population delays (i.e. time shifts) between stimulation sites (see Fig. 8B) can induce changes in the inter-population synaptic connectivity in a network of LIF neurons which are similar to the classical STDP profile [74]. Consistent with our results, the induction of significant plastic changes in inter-population connections was crucially affected by appropriate stimulus timing [74].

**Figure 8:**
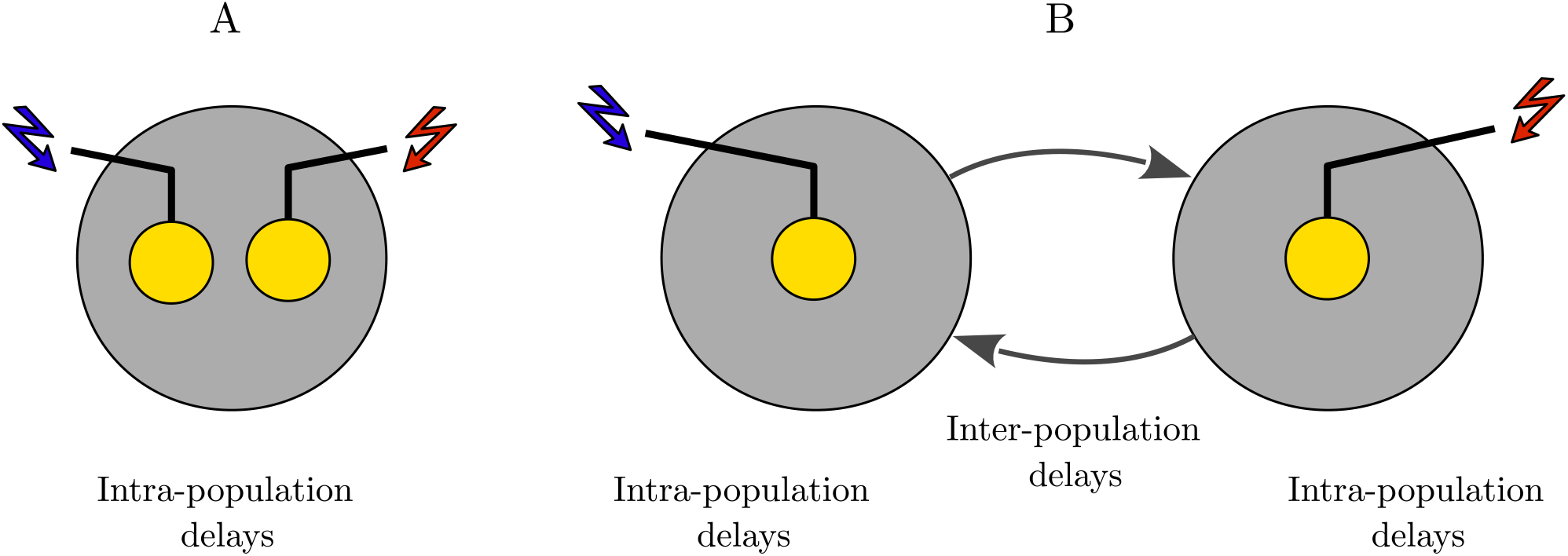
Illustrative representation of the time-shifted stimulation. (**A**) Schematically refers to a variety of bi-/multichannel stimulation techniques where the delays between different stimulation sites (yellow circles) are approximately the same. (**B**) Schematically refers to bi- /multichannel stimulation techniques with at least two qualitatively different delays, i.e., small delays within and long delays between stimulation sites.

Cortical stimulation may offer simpler ways to modulate abnormally synchronized activity which is less invasive in comparison to DBS surgery. However, cortical stimulation may also have its limitations. For instance, electrical stimulation can intrinsically evoke nonphysiological spatiotemporal patterns and complicated neural responses in cortical networks due to the transsynaptic activation of axons [73, 101]. Here, we used a simple and generic cortical model [60, 63, 76] to illustrate the critical role of the time shift in shaping long-lasting decoupling effects induced by stimulation. A simple model enables clear predictions based on a detailed investigation that is valid for a large parameter space, whereas complex and detailed computational models with many parameters may be difficult to study because of their high-dimensional phase space and the associated computational cost. Furthermore, we focused on the decoupling of the two populations achieved by the stimulation-induced reduction of the plastic synaptic connections between the populations and for simplicity assumed that the modules have only static synapses. However, assuming plastic synapses within each module may affect the presented results. In this case, the time-shifted stimulation should first decouple the two modules externally and, then, reduce the internal coupling of each module to ensure that long-lasting decoupling and desynchronizing effects can be achieved.

Introducing a time shift is a comparably simple procedure which does not require complex technical solutions, e.g., required for the detection of biomarkers, as in the case of demand-controlled stimulation [102]. By means of this maneuver (i.e., introducing the time shift), a variety of different bi-/multichannel stimulation protocols may have significantly different long-term effects. Our findings thus may have quite generic implications for different cortical stimulation strategies, e.g., through epicortical electrodes or intracortical electrodes. On the other hand, cortical networks may exhibit evoked or spontaneous collective oscillations in the absence of temporally structured stimuli, despite irregular and weakly correlated activity pattern of individual neurons, known as the normal state of the cortical activity [93, 94]. By the same token, physiological states in subcortical networks are characterized by uncorrelated neuronal activity as opposed to pathological states that are characterized by abnormally synchronized activity, for example, in PD [12]. This suggests that the results obtained for the simple cortical model may also be, in principle, applicable to basal ganglia models subjected to multichannel DBS protocols.

Interestingly, for the parameter ranges that we tested, qualitatively similar results were obtained for both the patterned delivery of bursts and trains of single stimulus pulses, implying that the time shift scheme works with burst stimulation as well as with continuous stimulation. This suggests that the reshaping of activity-connectivity patterns by time-shifted stimulus delivery may be a mechanism that applies to a larger class of stimulation patterns. This is encouraging for further test in more complex network models with more sophisticated intrinsic dynamics as well as in pre-clinical (animal) studies and clinical tests.

The role of the introduced time (phase) shift between stimulation signals delivered to the two populations is critical. Our results show that the time-shifted stimulation is more effective when the stimulation signals are almost in anti-phase (i.e., *π* phase lag or ∆*t ≈* 1*/*2*ν*), whereas the almost in-phase stimulation (with zero phase lag or ∆*t ≈* 0) failed to induce decoupling effects in our model. We used a typical STDP profile that is experimentally observed in cortex [6] and a reasonable range for cortico-cortical transmission delay [83], but in general, as predicted by our theoretical analysis, the effective time shift for stimulation deviates from anti-phase based on the inter-population transmission delay (see Fig. 2) and the shape of the STDP learning window (see Fig. 5). The phase-specificity of the decoupling effects in our model is consistence with previous experimental studies suggesting that in-phase cortical stimulation has a synchronizing effect on the neuronal activity in the target node, whereas anti-phase stimulation can cause desynchronizing effects [103]. Targeting multiple nodes of brain networks thus may enable stimulation strategies to improve stimulation outcome by phase manipulation between stimulation sites.

Finally, STDP-induced reorganization of cortical networks by stimulation has been addressed in a number of experimental works related to our computational results. For instance, paired stimulation [104], i.e., time-locked pairing of two stimulations delivered to two sites, allows to directly activate, e.g., cortico-cortical pathways connecting two areas, to study changes in inter-population connections through STDP [73,105]. Experimental findings in humans suggest that STDP-like changes in cortico-cortical inter-hemispheric connectivity crucially depend on the delay (time shift) between paired stimuli delivered to two sites (also see Fig. 8B) in the primary motor cortex [69,70], as well as paired stimulation protocols targeting intra-hemispheric connectivity with short-range [71, 106] and long-range [71, 72] pairing delays (also see Fig. 8A). Amazingly, bidirectional STDP-like changes in inter-hemispheric connections can be achieved by changing which hemisphere is stimulated first [107].

A recent *in vivo* study in awake, behaving monkeys demonstrated that STDP could be induced via time-shifted paired stimulation of two interacting cortical sites (namely, A and B) [73]. By assuming a fixed stimulation protocol and varying the delay (time shift) between stimuli, STDP-like changes in the strength of connections between the two sites were achieved which were similar to classical single-cell findings *in vitro* [4–6]. For instance, positive delays between paired stimuli (i.e., pre-before-post or A → B) led to a strengthening of inter-population synapses, whereas negative delays (i.e., post-before-pre or B → A) weakened the synaptic connections [73]. Moreover, small time shifts between stimuli (*<* 20 ms) induced significant synaptic changes, whereas greater time shifts (> 50 ms) fell outside the functional temporal window of STDP [73], similar to the classical STDP profile [5,6]. Interestingly, consistent with previous computational findings [74] and our results, a zero delay between paired stimulations did not induce significant changes in inter-population connections [73].

The success of computational studies [74] to reproduce experimental observations [73] suggests that simple cortical models incorporating STDP may be able to explain the neural mechanisms underlying cortical stimulation targeting synaptic plasticity. Our results thus may contribute to the further optimization of a variety of bi-/multichannel stimulation protocols aimed at the therapeutic reshaping of brain networks.

## Acknowledgements

PT gratefully acknowledges funding support by the John A. Blume Foundation and the Foundation for OCD Research (New Venture Fund 011665-2020-08-01). The funders had no role in study design, data collection and analysis, decision to publish, or preparation of the manuscript.

## CRediT Author Statement

**Mojtaba Madadi Asl:** Conceptualization, Methodology, Software, Formal analysis, Investigation, Writing - original draft, Writing - review & editing, Visualization. **Alireza Valizadeh:** Conceptualization, Methodology, Formal analysis, Writing - original draft, Writing - review & editing, Supervision, Project administration. **Peter A. Tass:** Conceptualization, Methodology, Formal analysis, Writing - original draft, Writing - review & editing, Supervision.

## Data Availability

All data generated or analyzed during this study are included in this published article and its supplementary material files.

## Appendix A. Supplementary Material

Supplementary material accompanies this article.

## References

[1] Song S, Abbott LF. Cortical development and remapping through spike timing-dependent plasticity. Neuron. 2001;32(2):339–350.

[2] Ramanathan D, Conner JM, Tuszynski MH. A form of motor cortical plasticity that correlates with recovery of function after brain injury. Proceedings of the National Academy of Sciences. 2006;103(30):11370–11375.

[3] Gerstner W, Kempter R, van Hemmen JL, Wagner H. A neuronal learning rule for sub-millisecond temporal coding. Nature. 1996;383(6595):76.

[4] Markram H, Lübke J, Frotscher M, Sakmann B. Regulation of synaptic efficacy by coincidence of postsynaptic APs and EPSPs. Science. 1997;275(5297):213–215.

[5] Bi GQ, Poo MM. Synaptic modifications in cultured hippocampal neurons: dependence on spike timing, synaptic strength, and postsynaptic cell type. Journal of Neuroscience. 1998;18(24):10464–10472.

[6] Froemke RC, Dan Y. Spike-timing-dependent synaptic modification induced by natural spike trains. Nature. 2002;416(6879):433–438.

[7] Morrison A, Aertsen A, Diesmann M. Spike-timing-dependent plasticity in balanced random networks. Neural Computation. 2007;19(6):1437–1467.

[8] Gilson M, Burkitt A, van Hemmen JL. STDP in recurrent neuronal networks. Frontiers in Computational Neuroscience. 2010;4:23.

[9] Madadi Asl M, Valizadeh A, Tass PA. Propagation delays determine neuronal activity and synaptic connectivity patterns emerging in plastic neuronal networks. Chaos. 2018;28(10):106308.

[10] Madadi Asl M, Valizadeh A, Tass PA. Dendritic and axonal propagation delays may shape neuronal networks with plastic synapses. Frontiers in Physiology. 2018;9:1849.

[11] Goldberg JA, Boraud T, Maraton S, Haber SN, Vaadia E, Bergman H. Enhanced synchrony among primary motor cortex neurons in the 1-methyl-4-phenyl-1, 2, 3, 6-tetrahydropyridine primate model of Parkinson’s disease. Journal of Neuroscience. 2002;22(11):4639–4653.

[12] Hammond C, Bergman H, Brown P. Pathological synchronization in Parkinson’s disease: networks, models and treatments. Trends in Neurosciences. 2007;30(7):357–364.

[13] Fan KY, Baufreton J, Surmeier DJ, Chan CS, Bevan MD. Proliferation of external globus pallidus-subthalamic nucleus synapses following degeneration of midbrain dopamine neurons. Journal of Neuroscience. 2012;32(40):13718–13728.

[14] Chu HY, Atherton JF, Wokosin D, Surmeier DJ, Bevan MD. Heterosynaptic regulation of external globus pallidus inputs to the subthalamic nucleus by the motor cortex. Neuron. 2015;85(2):364–376.

[15] Asadi A, Madadi Asl M, Vahabie AH, Valizadeh A. The origin of abnormal beta oscillations in the parkinsonian corticobasal ganglia circuits. Parkinson’s Disease. 2022;2022(7524066):1–13.

[16] Kane A, Hutchison WD, Hodaie M, Lozano AM, Dostrovsky JO. Enhanced synchronization of thalamic theta band local field potentials in patients with essential tremor. Experimental Neurology. 2009;217(1):171–176.

[17] Schnitzler A, Münks C, Butz M, Timmermann L, Gross J. Synchronized brain network associated with essential tremor as revealed by magnetoencephalography. Movement Disorders. 2009;24(11):1629–1635.

[18] Buijink AW, van der Stouwe AM, Broersma M, Sharifi S, Groot PF, Speelman JD, et al. Motor network disruption in essential tremor: a functional and effective connectivity study. Brain. 2015;138(10):2934–2947.

[19] Mormann F, Lehnertz K, David P, Elger CE. Mean phase coherence as a measure for phase synchronization and its application to the EEG of epilepsy patients. Physica D: Nonlinear Phenomena. 2000;144(3-4):358–369.

[20] da Silva FHL, Blanes W, Kalitzin SN, Parra J, Suffczynski P, Velis DN. Dynamical diseases of brain systems: different routes to epileptic seizures. IEEE Transactions on Biomedical Engineering. 2003;50(5):540–548.

[21] Chu Y, Jin X, Parada I, Pesic A, Stevens B, Barres B, et al. Enhanced synaptic connectivity and epilepsy in C1q knockout mice. Proceedings of the National Academy of Sciences. 2010;107(17):7975–7980.

[22] McGregor MM, Nelson AB. Circuit mechanisms of Parkinson’s disease. Neuron. 2019;101(6):1042–1056.

[23] Mallet N, Delgado L, Chazalon M, Miguelez C, Baufreton J. Cellular and synaptic dysfunctions in Parkinson’s disease: stepping out of the striatum. Cells. 2019;8(9):1005.

[24] Madadi Asl M, Vahabie AH, Valizadeh A, Tass PA. Spike-timing-dependent plasticity mediated by dopamine and its role in Parkinson’s disease pathophysiology. Frontiers in Network Physiology. 2022;2(817524):1–18.

[25] Madadi Asl M, Vahabie AH, Valizadeh A. Dopaminergic modulation of synaptic plasticity, its role in neuropsychiatric disorders, and its computational modeling. Basic and Clinical Neuroscience. 2019;10(1):1.

[26] Madadi Asl M, Asadi A, Enayati J, Valizadeh A. Inhibitory spike-timing-dependent plasticity can account for pathological strengthening of pallido-subthalamic synapses in Parkinson’s disease. Frontiers in Physiology. 2022;13(915626):1–13.

[27] Meissner W, Leblois A, Hansel D, Bioulac B, Gross CE, Benazzouz A, et al. Subthalamic high frequency stimulation resets subthalamic firing and reduces abnormal oscillations. Brain. 2005;128(10):2372–2382.

[28] Kühn AA, Kempf F, Brücke C, Doyle LG, Martinez-Torres I, Pogosyan A, et al. High-frequency stimulation of the subthalamic nucleus suppresses oscillatory β activity in patients with Parkinson’s disease in parallel with improvement in motor performance. Journal of Neuroscience. 2008;28(24):6165–6173.

[29] Temperli P, Ghika J, Villemure JG, Burkhard P, Bogousslavsky J, Vingerhoets F. How do parkinsonian signs return after discontinuation of subthalamic DBS? Neurology. 2003;60(1):78–81.

[30] Volkmann J. Deep brain stimulation for the treatment of Parkinson’s disease. Journal of Clinical Neurophysiology. 2004;21(1):6–17.

[31] Baizabal-Carvallo JF, Jankovic J. Movement disorders induced by deep brain stimulation. Parkinsonism & Related Disorders. 2016;25:1–9.

[32] Lozano AM, Lipsman N, Bergman H, Brown P, Chabardes S, Chang JW, et al. Deep brain stimulation: current challenges and future directions. Nature Reviews Neurology. 2019;15(3):148–160.

[33] Tass PA. Phase resetting in medicine and biology: stochastic modelling and data analysis. Springer Verlag, Berlin; 1999.

[34] Tass PA. Stochastic phase resetting: a theory for deep brain stimulation. Progress of Theoretical Physics Supplement. 2000;139:301–313.

[35] Rosenblum M, Pikovsky A. Delayed feedback control of collective synchrony: An approach to suppression of pathological brain rhythms. Physical Review E. 2004;70(4):041904.

[36] Rosenblum MG, Pikovsky AS. Controlling synchronization in an ensemble of globally coupled oscillators. Physical Review Letters. 2004;92(11):114102.

[37] Popovych OV, Hauptmann C, Tass PA. Effective desynchronization by nonlinear delayed feedback. Physical Review Letters. 2005;94(16):164102.

[38] Hauptmann C, Popovych O, Tass PA. Delayed feedback control of synchronization in locally coupled neuronal networks. Neurocomputing. 2005;65:759–767.

[39] Pyragas K, Popovych O, Tass P. Controlling synchrony in oscillatory networks with a separate stimulation-registration setup. Europhysics Letters. 2007;80(4):40002.

[40] Ratas I, Pyragas K. Controlling synchrony in oscillatory networks via an act-and-wait algorithm. Physical Review E. 2014;90(3):032914.

[41] Kuncel AM, Grill WM. Selection of stimulus parameters for deep brain stimulation. Clinical Neurophysiology. 2004;115(11):2431–2441.

[42] Harnack D, Winter C, Meissner W, Reum T, Kupsch A, Morgenstern R. The effects of electrode material, charge density and stimulation duration on the safety of high-frequency stimulation of the subthalamic nucleus in rats. Journal of Neuroscience Methods. 2004;138(1-2):207–216.

[43] Merrill DR, Bikson M, Jefferys JG. Electrical stimulation of excitable tissue: design of efficacious and safe protocols. Journal of Neuroscience Methods. 2005;141(2):171–198.

[44] Popovych OV, Lysyansky B, Rosenblum M, Pikovsky A, Tass PA. Pulsatile desynchronizing delayed feedback for closed-loop deep brain stimulation. PloS ONE. 2017;12(3):e0173363.

[45] Popovych OV, Lysyansky B, Tass PA. Closed-loop deep brain stimulation by pulsatile delayed feedback with increased gap between pulse phases. Scientific Reports. 2017;7(1):1– 14.

[46] Popovych OV, Tass PA. Adaptive delivery of continuous and delayed feedback deep brain stimulation-a computational study. Scientific Reports. 2019;9(1):1–17.

[47] Grill WM. Temporal pattern of electrical stimulation is a new dimension of therapeutic innovation. Current Opinion in Biomedical Engineering. 2018;8:1–6.

[48] Khaledi-Nasab A, Kromer JA, Tass PA. Long-lasting desynchronization of plastic neural networks by random reset stimulation. Frontiers in Physiology. 2020;11:622620.

[49] Khaledi-Nasab A, Kromer JA, Tass PA. Long-lasting desynchronization of plastic neuronal networks by double-random coordinated reset stimulation. Frontiers in Network Physiology. 2022;2:864859.

[50] Tass PA. A model of desynchronizing deep brain stimulation with a demand-controlled coordinated reset of neural subpopulations. Biological Cybernetics. 2003;89(2):81–88.

[51] Tass PA, Majtanik M. Long-term anti-kindling effects of desynchronizing brain stimulation: a theoretical study. Biological Cybernetics. 2006;94(1):58–66.

[52] Hauptmann C, Tass PA. Cumulative and after-effects of short and weak coordinated reset stimulation: a modeling study. Journal of Neural Engineering. 2009;6(1):016004.

[53] Tass PA, Qin L, Hauptmann C, Dovero S, Bezard E, Boraud T, et al. Coordinated reset has sustained aftereffects in Parkinsonian monkeys. Annals of Neurology. 2012;72(5):816– 820.

[54] Adamchic I, Hauptmann C, Barnikol UB, Pawelczyk N, Popovych O, Barnikol TT, et al. Coordinated reset neuromodulation for Parkinson’s disease: proof-of-concept study. Movement Disorders. 2014;29(13):1679–1684.

[55] Wang J, Nebeck S, Muralidharan A, Johnson MD, Vitek JL, Baker KB. Coordinated reset deep brain stimulation of subthalamic nucleus produces long-lasting, dose-dependent motor improvements in the 1-methyl-4-phenyl-1, 2, 3, 6-tetrahydropyridine non-human primate model of parkinsonism. Brain Stimulation. 2016;9(4):609–617.

[56] Bore JC, Campbell BA, Cho H, Pucci F, Gopalakrishnan R, Machado AG, et al. Long-lasting effects of subthalamic nucleus coordinated reset deep brain stimulation in the non-human primate model of parkinsonism: A case report. Brain Stimulation. 2022;.

[57] Wang J, Fergus SP, Johnson LA, Nebeck SD, Zhang J, Kulkarni S, et al. Shuffling Improves the Acute and Carryover Effect of Subthalamic Coordinated Reset Deep Brain Stimulation. Frontiers in Neurology. 2022;13:716046–716046.

[58] Manos T, Zeitler M, Tass PA. How stimulation frequency and intensity impact on the long-lasting effects of coordinated reset stimulation. PLoS Computational Biology. 2018;14(5):e1006113.

[59] Kromer JA, Tass PA. Long-lasting desynchronization by decoupling stimulation. Physical Review Research. 2020;2(3):033101.

[60] Diesmann M, Gewaltig MO, Aertsen A. Stable propagation of synchronous spiking in cortical neural networks. Nature. 1999;402(6761):529–533.

[61] Akam T, Kullmann DM. Oscillations and filtering networks support flexible routing of information. Neuron. 2010;67(2):308–320.

[62] Hahn G, Bujan AF, Frégnac Y, Aertsen A, Kumar A. Communication through resonance in spiking neuronal networks. PLoS Computational Biology. 2014;10(8):e1003811.

[63] Rezaei H, Aertsen A, Kumar A, Valizadeh A. Facilitating the propagation of spiking activity in feedforward networks by including feedback. PLOS Computational Biology. 2020;16(8):e1008033.

[64] Barardi A, Sancristóbal B, Garcia-Ojalvo J. Phase-coherence transitions and communication in the gamma range between delay-coupled neuronal populations. PLoS Computational Biology. 2014;10(7):e1003723.

[65] Viaro R, Morari M, Franchi G. Progressive motor cortex functional reorganization following 6-hydroxydopamine lesioning in rats. Journal of Neuroscience. 2011;31(12):4544–4554.

[66] Underwood CF, Parr-Brownlie LC. Primary motor cortex in Parkinson’s disease: Functional changes and opportunities for neurostimulation. Neurobiology of Disease. 2021;147:105159.

[67] Drouot X, Oshino S, Jarraya B, Besret L, Kishima H, Remy P, et al. Functional recovery in a primate model of Parkinson’s disease following motor cortex stimulation. Neuron. 2004;44(5):769–778.

[68] Fregni F, Boggio PS, Santos MC, Lima M, Vieira AL, Rigonatti SP, et al. Noninvasive cortical stimulation with transcranial direct current stimulation in Parkinson’s disease. Movement Disorders. 2006;21(10):1693–1702.

[69] Rizzo V, Siebner H, Morgante F, Mastroeni C, Girlanda P, Quartarone A. Paired associative stimulation of left and right human motor cortex shapes interhemispheric motor inhibition based on a Hebbian mechanism. Cerebral Cortex. 2009;19(4):907–915.

[70] Koganemaru S, Mima T, Nakatsuka M, Ueki Y, Fukuyama H, Domen K. Human motor associative plasticity induced by paired bihemispheric stimulation. The Journal of physiology. 2009;587(19):4629–4644.

[71] Buch ER, Johnen VM, Nelissen N, O’Shea J, Rushworth MF. Noninvasive associative plasticity induction in a corticocortical pathway of the human brain. Journal of Neuroscience. 2011;31(48):17669–17679.

[72] Koch G, Ponzo V, Di Lorenzo F, Caltagirone C, Veniero D. Hebbian and anti-Hebbian spike-timing-dependent plasticity of human cortico-cortical connections. Journal of Neuroscience. 2013;33(23):9725–9733.

[73] Seeman SC, Mogen BJ, Fetz EE, Perlmutter SI. Paired stimulation for spike-timing-dependent plasticity in primate sensorimotor cortex. Journal of Neuroscience. 2017;37(7):1935–1949.

[74] Shupe L, Fetz E. An integrate-and-fire spiking neural network model simulating artificially induced cortical plasticity. eNeuro. 2021;8(2).

[75] Burkitt AN. A review of the integrate-and-fire neuron model: I. Homogeneous synaptic input. Biological Cybernetics. 2006;95(1):1–19.

[76] Sadeh S, Clopath C. Excitatory-inhibitory balance modulates the formation and dynamics of neuronal assemblies in cortical networks. Science Advances. 2021;7(45):eabg8411.

[77] Buzsáki G, Mizuseki K. The log-dynamic brain: how skewed distributions affect network operations. Nature Reviews Neuroscience. 2014;15(4):264–278.

[78] Brunel N, Wang XJ. What determines the frequency of fast network oscillations with irregular neural discharges? I. Synaptic dynamics and excitation-inhibition balance. Journal of Neurophysiology. 2003;90(1):415–430.

[79] Madadi Asl M, Valizadeh A, Tass PA. Dendritic and axonal propagation delays determine emergent structures of neuronal networks with plastic synapses. Scientific Reports. 2017;7:39682.

[80] Kumar A, Rotter S, Aertsen A. Conditions for propagating synchronous spiking and asynchronous firing rates in a cortical network model. Journal of Neuroscience. 2008;28(20):5268–5280.

[81] Roohi N, Valizadeh A. Role of Interaction Delays in the Synchronization of Inhibitory Networks. Neural Computation. 2022;34(6):1425–1447.

[82] Madadi Asl M, Valizadeh A, Tass PA. Delay-induced multistability and loop formation in neuronal networks with spike-timing-dependent plasticity. Scientific Reports. 2018;8:12068.

[83] Lemaréchal JD, Jedynak M, Trebaul L, Boyer A, Tadel F, Bhattacharjee M, et al. A brain atlas of axonal and synaptic delays based on modelling of cortico-cortical evoked potentials. Brain. 2021;.

[84] Aoki T, Aoyagi T. Co-evolution of phases and connection strengths in a network of phase oscillators. Physical Review Letters. 2009;102(3):034101.

[85] Babadi B, Abbott LF. Pairwise analysis can account for network structures arising from spike-timing dependent plasticity. PLoS Computational Biology. 2013;9(2):e1002906.

[86] Schmalz J, Kumar G. Controlling synchronization of spiking neuronal networks by harnessing synaptic plasticity. Frontiers in Computational Neuroscience. 2019;13:61.

[87] Oswal A, Brown P, Litvak V. Synchronized neural oscillations and the pathophysiology of Parkinson’s disease. Current Opinion in Neurology. 2013;26(6):662–670.

[88] Stoelzel CR, Bereshpolova Y, Alonso JM, Swadlow HA. Axonal conduction delays, brain state, and corticogeniculate communication. Journal of Neuroscience. 2017;37(26):6342– 6358.

[89] Kozloski J, Cecchi GA. A theory of loop formation and elimination by spike timing-dependent plasticity. Frontiers in Neural Circuits. 2010;4:7.

[90] Knoblauch A, Hauser F, Gewaltig MO, Körner E, Palm G. Does spike-timing-dependent synaptic plasticity couple or decouple neurons firing in synchrony? Frontiers in Computational Neuroscience. 2012;6(55):55.

[91] Lubenov EV, Siapas AG. Decoupling through synchrony in neuronal circuits with propagation delays. Neuron. 2008;58(1):118–131.

[92] Popovych OV, Hauptmann C, Tass PA. Desynchronization and decoupling of interacting oscillators by nonlinear delayed feedback. International Journal of Bifurcation and Chaos. 2006;16(07):1977–1987.

[93] Arieli A, Sterkin A, Grinvald A, Aertsen A. Dynamics of ongoing activity: explanation of the large variability in evoked cortical responses. Science. 1996;273(5283):1868–1871.

[94] Renart A, De La Rocha J, Bartho P, Hollender L, Parga N, Reyes A, et al. The asynchronous state in cortical circuits. Science. 2010;327(5965):587–590.

[95] Tass PA, Hauptmann C. Therapeutic modulation of synaptic connectivity with desynchronizing brain stimulation. International Journal of Psychophysiology. 2007;64(1):53– 61.

[96] Song S, Miller KD, Abbott LF. Competitive Hebbian learning through spike-timing-dependent synaptic plasticity. Nature Neuroscience. 2000;3(9):919–926.

[97] Popovych OV, Tass PA. Desynchronizing electrical and sensory coordinated reset neuromodulation. Frontiers in Human Neuroscience. 2012;6(58):58.

[98] Kromer JA, Khaledi-Nasab A, Tass PA. Impact of number of stimulation sites on long-lasting desynchronization effects of coordinated reset stimulation. Chaos. 2020;30(8):083134.

[99] Khaledi-Nasab A, Kromer JA, Tass PA. Long-lasting desynchronization effects of coordinated reset stimulation improved by random jitters. Frontiers in Physiology. 2021;(719680):1446.

[100] Khaledi-Nasab A, Kromer JA, Tass PA. Long-lasting desynchronization of plastic neural networks by random reset stimulation. Frontiers in Physiology. 2021;(622620):1843.

[101] Histed MH, Bonin V, Reid RC. Direct activation of sparse, distributed populations of cortical neurons by electrical microstimulation. Neuron. 2009;63(4):508–522.

[102] Hoang KB, Cassar IR, Grill WM, Turner DA. Biomarkers and stimulation algorithms for adaptive brain stimulation. Frontiers in Neuroscience. 2017;11:564.

[103] Alagapan S, Riddle J, Huang WA, Hadar E, Shin HW, Fröhlich F. Network-targeted, multi-site direct cortical stimulation enhances working memory by modulating phase lag of low-frequency oscillations. Cell Reports. 2019;29(9):2590–2598.

[104] Stefan K, Kunesch E, Cohen LG, Benecke R, Classen J. Induction of plasticity in the human motor cortex by paired associative stimulation. Brain. 2000;123(3):572–584.

[105] Santarnecchi E, Momi D, Sprugnoli G, Neri F, Pascual-Leone A, Rossi A, et al. Modulation of network-to-network connectivity via spike-timing-dependent noninvasive brain stimulation. Human Brain Mapping. 2018;39(12):4870–4883.

[106] Arai N, Müller-Dahlhaus F, Murakami T, Bliem B, Lu MK, Ugawa Y, et al. State-dependent and timing-dependent bidirectional associative plasticity in the human SMA-M1 network. Journal of Neuroscience. 2011;31(43):15376–15383.

[107] Zibman S, Daniel E, Alyagon U, Etkin A, Zangen A. Interhemispheric cortico-cortical paired associative stimulation of the prefrontal cortex jointly modulates frontal asymmetry and emotional reactivity. Brain Stimulation. 2019;12(1):139–147.

