## Supplementary Material for "Decoupling of interacting neuronal populations by time-shifted stimulation through spike-timing-dependent plasticity"

### Two-Neuron Motif in the Absence of Delay

The numerical results shown in Fig. S1A for an exemplary set of parameters (marked by point b in Fig. 2A) confirm the theoretical predictions in Fig. 2A when delays are not considered in the model. In this case, the spontaneous firing of the neurons led to unidirectional potentiation of the synapses before stimulation onset (point a in Fig. 2A). Stimulation of the two neurons with a proper time shift ($\Delta t$) and frequency ($\nu = 1/T$) corresponding to parameters marked by point b: $(\Delta t, T) = (15, 30)$ ms in Fig. 2A, shifted the dynamics of the motif towards the depression regime (point c in Fig. 2A), resulting in the depression of both synapses.

### Bidirectionally Connected Populations in the Absence of Delay

Raster plot of the firing of the neurons in two populations is shown in Fig. S2A1 and A2 when delays were ignored in the model. Before stimulation, the mean pairwise correlation between the firing of the neurons in each module is high and the neurons' firing show a high temporal regularity (Fig. S2B and C, grey). The distribution of phase difference between the oscillations of the two modules and the frequencies of the network oscillations (before stimulation onset) shown in Fig. S2D (left) and E (grey) indicate that this regime is expected to shape a unidirectional connectivity (like point a in Fig. 2A in the two-neuron motif). This is verified by the results shown in Fig. S2F and G (grey) where the time course of the mean inter-population coupling and the distribution of the synaptic strengths are represented.

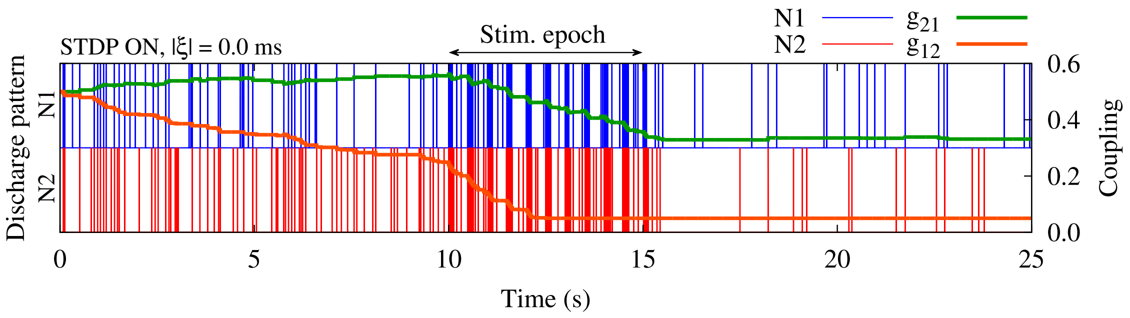


Figure S1: **Suppression of the synaptic strengths between two neurons by time-shifted stimulation.** Time course of neuronal activity (N1/N2) and the synaptic strengths ($g_{21}/g_{12}$) are shown for two neurons with STDP parameters $A_+ = 0.008$, $A_- = 0.005$, $\tau_+ = 10$ ms and $\tau_- = 20$ ms, and $|\xi| = 0.0$ ms.

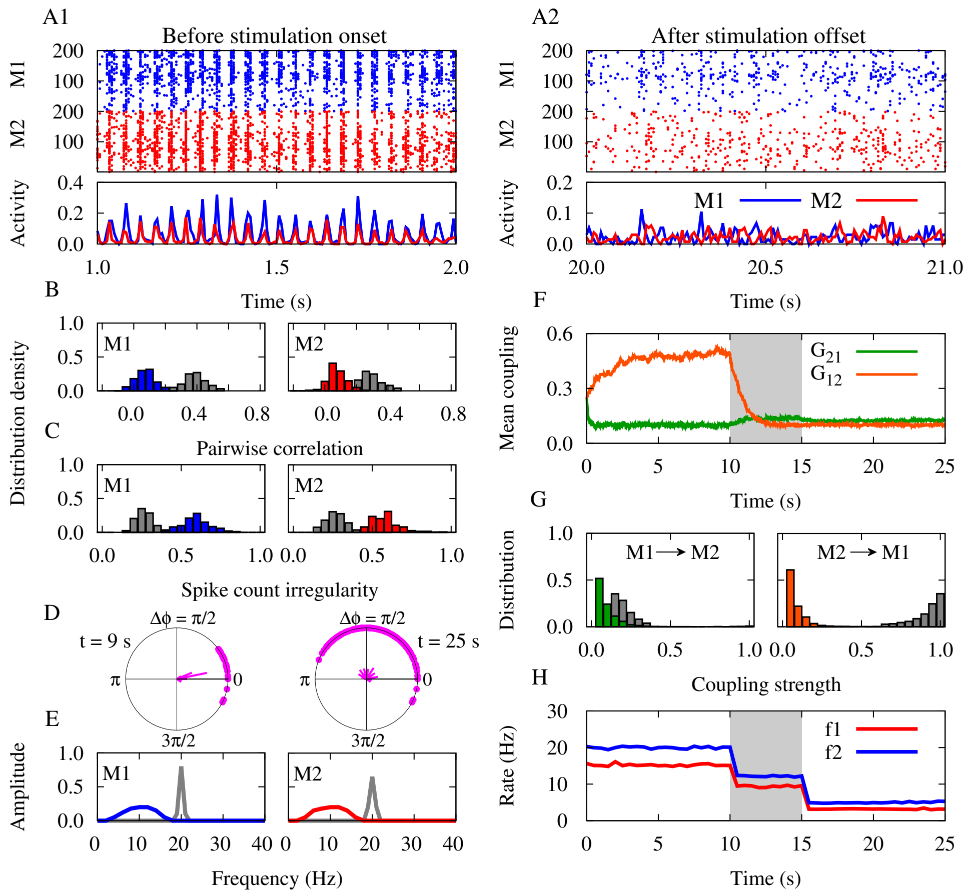


Figure S2: **Decoupling by time-shifted stimulation in the network.** (**A1,A2**) Raster plots and population activities are shown for the modules (M1/M2) before/after stimulation on/offset. (**B,C**) Distribution of the pairwise correlation and spike count irregularity of each module before (grey) and after (colored) stimulation. (**D**) Snapshot of phase lags ($\Delta\phi$) between synchronous discharges from the two modules before ($t = 9\,$s, left) and after ($t = 25\,$s, right) stimulation. (**E**) Fourier transform frequency of the population activity of each module before (grey) and after (colored) stimulation. (**F**) Time course of the inter-population mean coupling ($G_{21}/G_{12}$). (**G**) Distribution of the inter-population synaptic strengths before (grey) and after (colored) stimulation. (**H**) Time course of the firing rate of neurons ($f1/f2$) in each module. The modules were stimulated with time shift $\Delta t = 15\,$ms and frequency $\nu = 1/30\,$Hz (point b in Fig. 2A) for the duration of $T_{\text{stim}} = 5\,$s (highlighted area in F and H). STDP parameters were $A_+ = 0.008$, $A_- = 0.005$, $\tau_+ = 10\,$ms and $\tau_- = 20\,$ms.

We then stimulated the two populations with the time shift $\Delta t = 15\,$ms and frequency $\nu = 1/30\,$Hz corresponding to parameters shown by point b in Fig. 2A. The strong connections from module $2 \longrightarrow 1$ are suppressed while no significant change is seen in the weak connections in the reverse direction (see Fig. S2F and G, colored). The decoupling of the two populations led to the unlearning of pathological dynamics so that strong oscillations disappeared and only weak oscillations remained (Fig. S2E, colored). The firing rate of neurons decreased considerably (Fig. S2H) where the neurons fire in an irregular (Fig. S2C, colored) and sparse manner, i.e., with a much lower rate compared to the frequency of oscillations. The pairwise correlation between the spiking of the neurons decreases (Fig. S2B, colored) and the phase difference between the spiking of the neurons in the two modules takes a wider distribution (Fig. S2D, right).
